# The Immune landscape of pediatric solid tumors

**DOI:** 10.1101/2021.05.04.442503

**Authors:** Shimaa Sherif, Jessica Roelands, William Mifsud, Eiman Ahmed, Borbala Mifsud, Davide Bedognetti, Wouter Hendrickx

**Affiliations:** College of Health and Life Sciences (CHLS), Hamad Bin Khalifa University (HBKU), Doha, Qatar; Pediatric Cancer Omics Lab, Cancer Program, Research Branch, Sidra Medicine, Doha, Qatar; Department of Pathology, Leiden University Medical Center, Leiden, The Netherlands; Anatomical Pathology department, Sidra Medicine, Doha, Qatar; Cancer Immunogenetics Lab, Cancer Program, Research Branch, Sidra Medicine, Doha, Qatar

**Keywords:** Pan-cancer, Immunologic Constant of Rejection, Prognosis, Tumor intrinsic pathway enrichment, Immune subtypes, Cancer Immunotherapy

## Abstract

**Background:** Immunotherapy is quickly coming to the forefront of cancer treatment; however, the implementation of immunotherapy in solid pediatric cancers, which classically display a low mutational load, is hindered by insufficient understanding of the determinants of cancer immune responsiveness in children. In order to better understand tumor-host interplay, we sought to characterize solid pediatric cancers based on immunological parameters using analytes extracted from gene expression data.

**Methods:** We used RNAseq data from the publicly available TARGET studies for five pediatric solid tumor types (408 patients): Wilms tumor (WT), neuroblastoma (NBL), osteosarcoma (OS), clear cell sarcoma of the kidney (CCSK) and rhabdoid tumor of the kidney (RT). We assessed the performance of previously identified immune signatures like the Immunologic Constant of Rejection (ICR), which captures an active Th1/cytotoxic response associated with favorable prognosis and responsiveness to immunotherapy. We also performed gene set enrichment analysis (ssGSEA) and clustering, using more than 100 immune signatures to define immune subtypes in pediatric tumors and compared the overall survival across subtypes. The expression of immune checkpoints and enrichment of oncogenic pathways were also assessed across the immune subtypes.

**Results:** The five tumor types showed distinct ICR score distributions. A higher ICR score was associated with better survival in OS and NBL-HR-MYCN_NA, but with poorer survival in WT. The clustering of immune signatures revealed the same five principal modules observed in adult solid tumors: Wound Healing, TGF-B signaling, IFN-G signaling, Macrophages, and Lymphocytes. These modules clustered pediatric patients into six immune subtypes (S1-S6) with distinct survival outcomes. The S2 cluster showed the best overall survival and was characterized by low enrichment of the wound healing signature, high Th1, low Th2. Conversely, cluster S4 showed the worst survival and highest enrichment of wound healing signature, low Th1, and high Th2. Furthermore, the upregulation of the WNT/Beta-catenin pathway is associated with unfavorable outcomes and lack of T-cell infiltration in OS.

**Conclusions:** We demonstrated that extracranial solid pediatric tumors could be classified according to their immune disposition, unveiling similarity with adults’ tumors. Immunological parameters might be explored to refine diagnostic and prognostic biomarkers and to identify potential immune-responsive tumors.

## Background

Cancer is one of the leading causes of death in children worldwide, and the recorded incidence tends to raise with time (Steliarova-Foucher et al. 2017). The overall incidence rates of childhood cancer vary between 50 and 200 per million children across the world (Steliarova-Foucher et al. 2017). The most common categories of childhood cancers include leukemias, brain tumors, lymphomas, neuroblastoma and Wilms tumor (Gelband et al. 2015). Solid tumors comprise almost half of the cancer cases (J. A. Lee 2018), and neuroblastoma, nephroblastoma are tumor types that occur almost exclusively in children(Quintero Escobar et al. 2019).

Although major progress has been made in the treatment of pediatric cancers since the 1960s, and despite intensification of treatment, contemporary progress has been limited or absent (Adamson 2015), (Dome, Perlman, and Graf 2014). For those where the disease relapses, survival remains poor and for patients presenting with non-invasive tumors, surgical intervention can cause major long-term consequences. In addition, many children with high-risk cancers develop severe, life-threatening, or fatal toxicity during their medical treatment (Crist et al. 1995).

It is now well established that the immune system has a substantial role in controlling cancer growth and stemming progression, and immunotherapy is quickly coming to the forefront of cancer treatment. Immunotherapy has rapidly changed the therapeutic landscape and prognosis for many hematologic malignancies (Hallek et al. 2010)(Burger et al. 2015)(Moreno et al. 2019) and adult solid tumors (Keilson et al. 2021). However, immunotherapy response depends on the tumor’s immune phenotype, therefore, it is important to characterize the tumor’s immune infiltration. Immune mediated tumor rejection can be estimated using gene expression signatures that stratify tumors into immune active (hot) and immune silent (cold) phenotypes. One such immune signature, which is prognostic in adult tumors, is the Immunologic Constant of Rejection (ICR). The ICR consists of 20 genes that reflect activation of Th-1-signaling (IFNG, TXB21, CD8B, CD8A, IL12B, STAT1, and IRF1), expression of CXCR3/CCR5 chemokine ligands (CXCL9, CXCL10, and CCL5), cytotoxic effector molecules (GNLY, PRF1, GZMA, GZMB, and GZMH) and compensatory immune regulators (CD274/PD-L1, PDCD1, CTLA4, FOXP3 and IDO1). Furthermore, T helper 1 (Th1) immune active phenotype is associated with responsiveness to immunotherapy and favorable prognosis in most but not all tumor types (Thorsson et al. 2018; Galon et al. 2013; Roelands et al. 2020).

These tumor-type-specific differences might be due to differences in the underlying oncogenic dysregulations. For instance, we recently demonstrated that the prognostic value of Th1 immune infiltration is dependent on the activation status of specific oncogenic processes such as proliferation and *TGF-B* signaling, even within the same cancer type (Bertucci et al. 2018; Roelands et al. 2019). Therefore, it is important to assess immune phenotypes in relation to activated oncogenic pathways in order to understand the differences in prognosis and response.

Despite the successes of immunotherapy in adult cancers, immunotherapy for pediatric solid tumors remains in the early stages of development, and significant clinical benefit is yet to be realized, except for the anti-GD2 antibody that is used for treating neuroblastoma (Yu et al. 2010)(Ladenstein et al. 2018). Several challenges prevent the adoption of immunotherapeutic strategies that have been successful in other settings. A relatively stable cancer genome with a low mutational burden results in limited neoepitope expression, which decreases T-cell infiltration to the tumor microenvironment, rendering the inhibition of exhaustion checkpoints like PD1/PDL1 and CTLA-4 less effective (Rizvi et al. 2015)(Buchbinder and Desai 2016). Moreover, the efficiency of immunotherapy depends on the specific molecular subtype of a given tumor, but identification of molecular subtypes is more difficult in pediatric tumors, since they are much rarer than their adult counterparts.

The development of immunotherapeutic strategies for solid pediatric cancers requires better understanding of the biology of the tumor immune microenvironment and identification of immune checkpoints that could be targeted by current or future checkpoint inhibitors (Roelands et al. 2020; Hendrickx et al. 2017).

In this study we analyzed transcriptomic data to characterize the immune infiltrate of solid pediatric tumors to define immune phenotypes in these tumors.

## Methods

All analysis was done in R version 3.6.1, software names are R packages unless stated otherwise.

### Data acquisition and normalization

RNA-seq data for 5 pediatric tumors: Wilms tumor (WT), neuroblastoma (NBL), osteosarcoma (OS), rhabdoid tumor (RT) and clear cell sarcoma of the kidney (CCSK) from the TARGET pediatric dataset, which is published on the GDC portal website, were downloaded and processed using TCGAbiolinks (v. 2.14.1)(Colaprico et al. 2016). Gene symbols were converted to official HGNC symbols using TCGAbiolinks, genes without symbol or gene information were excluded and this resulted in a pan-cancer expression matrix with 20,155 genes. Metastatic tumor, recurrent primary tumor or blood derived samples were excluded and a single primary tumor (TP) sample was analyzed for each patient.

RNA-Seq gene counts were normalized using the TCGAanalyze_Normalization function from TCGAbiolinks, including within-lane normalization procedures to adjust for GC-content effect, between-lane normalization procedures to adjust for distributional differences between lanes as sequencing depth and quantile normalized using TCGAbiolinks. After normalization, the pan-cancer matrix was split per cancer type and log2 transformed and the clinical data of the TARGET study were obtained from the GDC portal.

### ICR classification

The gene expression data of the ICR signature used to cluster the patients from each cancer type and pan-cancer using the ConsensusClusterPlus (Wilkerson and Hayes 2010) (v.1.42.0) with the following parameters: 5000 repeats, a maximum of six clusters and agglomerative hierarchical clustering with Ward.D2 criterion. The optimal number of clusters was determined based on the Calinski-Harabasz index.

The three obtained clusters were annotated as ‘ICR High’, ‘ICR Medium’ and ‘ICR Low’, where ‘ICR High’ showed the highest expression of ICR genes and ‘ICR Low’ the lowest. ICR score was calculated for each sample, defined as the mean of the normalized, log2-transformed gene expression values of the ICR genes. Heatmaps were drawn using the ComplexHeatmap package (Gu, Eils, and Schlesner 2016).

T-distributed stochastic neighbor embedding (t-SNE) plot was used as a dimension reduction technique on the complete RNA expression matrix using Rtsne (Krijthe 2015) (v. 0.15) (settings perplexity=15, theta=0.5). t-SNE plots were annotated for cancer types and ICR clusters.

### Survival analysis

Clinical files contain survival data and clinical parameters such as age at diagnosis, tissue type, vital status, disease stage and disease metastasis, amongst others. For the overall survival analysis, we used the time to death and time to last follow up, vital status. For the event free survival, we considered; 1) relapse, 2) progression, 3) second malignant neoplasm death, and 4) death without remission, as events. We performed Kaplan-Meier survival analyses using the ggsurv function from survminer (v. 0.4.8). Patients with less than 1 day of follow-up were excluded and survival data were censored after a follow-up duration of 10 years. Hazards Ratio (HR) between ICR Low and ICR High groups or between the six immune subtypes and the corresponding p-values were calculated using X^2^ test. Confidence intervals (97.5% CIs) were defined using survival (v. 2.41–3).

Cox proportional hazards regression analysis was performed using Survival and visualized as a forest plot. Cancer types were added as a factor in the multivariate analysis. We applied the cox.zph function, to test the proportional hazards assumption (PHA). The same method was used to correct for the clinical parameters that contribute to the survival across immune subtypes in the High-risk *MYCN* not-amplified NBL cohort. These clinical parameters are the (Mitosis-Karyorrhexis Index) MKI (High, Intermediate and Low), Ploidy (diploid and hyperploid), Age group (0-18m,18m-5y and above 5y). Forest plots were generated using forestplot (v.1.7.2).

### Immune cell subpopulation and oncogenic pathway enrichment analysis

To determine the enrichment of particular gene sets, that reflect either immune cell types or certain oncogenic pathways, we performed single sample gene set enrichment analysis (ssGSEA) on the log2 transformed, normalized gene expression data. Immune cell-specific signatures were used as described in Bindea et al. (Bindea et al. 2013) with slight modification. The dendritic cell (DC) signature was replaced by immature dendritic cells (iDC), plasmacytoid dendritic cells (pDC), myeloid dendritic cells (mDC) and DC. Additionally, the regulatory T-cell (Treg) signature was used as described in Angelova et al. (Angelova et al. 2015). Gene sets that reflect specific tumor-related pathways were selected from multiple sources as described in detail in (Salerno et al. 2016)(Hendrickx et al. 2017)(Lu et al. 2017), and gene sets reflecting cancer related immune signatures were used as previously described by Thorsson et al. (Thorsson et al. 2018). The association between continuous gene set enrichment scores and survival was calculated as described above. Differences between the HRs of signatures were illustrated in a heatmap (ComplexHeatmap (v. 2.2.0)) and the p values were calculated by the cox formula. Signatures with a p value > 0.1 across all tumors were excluded.

### Comprehensive pediatric immune subtypes

ssGSEA was performed using 105 of the 108 previously described immune signatures (Thorsson et al. 2018). Three signatures were excluded from the analysis due to missing gene expression information. Spearman correlation between the resulting enrichment scores was calculated and visualized using corrplot. Signature modules were identified visually and then patients were clustered according to the ssGSEA enrichment of the 5 signatures representing the identified modules, previously identified by Thorsson etal. Sample clustering was carried out using k-means clustering (km = 6, repeats = 10000), using ComplexHeatmap. The gap statistics was used to calculate the optimal number of clusters.

Stacked bar chart from ggplot2 (v. 3.3.3). was used to show the percentage of each cancer type in the immune subtypes and the percentage of each immune subtype within each cancer type.

Density plots from ggplot2 were used to show the median of enrichment scores of selected immune signatures from the 105 signatures (Thorsson et al. 2018), in addition to the log2 values of *HLA-1* and *HLA-2* from the filtered normalized RNASeq matrix.

### Gene expression correlation

Correlation matrices of ICR genes expression were generated by calculating the Pearson correlation of the ICR genes’ expression within cancer types and pan-cancer using the corrplot package, CCSK was excluded from this correlation analysis because of the small sample size (n=13).

Spearman correlation was performed on the enrichment matrix of 105 tumor immune expression signatures (Thorsson et al. 2018) and plotted using corrplot (v. 0.84).

Correlation matrices of NK cells / CD8T enrichment scores and the enrichment score (ES) of selected oncogenic pathways were calculated using Pearson correlation and plotted by ComplexHeatmap.

### Immune checkpoints expression

We used a list of immune checkpoints, divided into activating and inhibitory. The median values of the log2 transformation of the normalized gene expression counts of these genes were used and plotted by ComplexHeatmap.

### CIBERSORTx immune cells fractions

In order to compare the immune cell fractions between different immune subtypes, we analyzed the normalized gene expression data of the 408 pediatric samples using the CIBERSORTx website. The relative proportions of 22 immune cell types within the leukocyte compartment (LM22) were estimated. Cell fractions were visualized in barcharts and boxplots using ggplot2.

We summed the proportions of related immune cells together in ‘Aggregates’ to facilitate comparisons (Thorsson et al. 2018).

Lymphocytes are the sum of B cells naive, B cells memory, T cells CD4 naive, T cells CD4 memory resting, T cells CD4 memory activated and T cells follicular helper, T cells regulatory, Tregs, T cells gamma delta, T cells CD8, NK cells resting, NK cells activate and Plasma cells fractions. Macrophages are the sum of Monocytes, Macrophages M0, Macrophages M1 and Macrophages M2 fractions. Dendritic cells are the sum of Dendritic cells resting and Dendritic cells activated fractions. Mast cells are the sum of Mast cells resting and Mast cells activated fractions.

## Results

### Prognostic value of ICR is different in pediatric tumors

We analysed the expression profiles of patient samples from five distinct solid pediatric cancer types: WT, NBL, OS, RT and CCSK from TARGET. After exclusion of the following patients: 20 OS patients, who were older than 18 years-old; one NBL patient, who did not have MYCN status information; one RT patient, whose sample clustered with NBL samples based on the whole transcriptome; we analyzed 408 patient samples (WT (n = 118), NBL (n = 150), OS (n = 68), RT (n= 59), CCSK (n= 13)). The NBL cohort was separated into three groups based on the COG risk group and the *MYCN* gene amplification status: High-risk MYCN amplified (NBL-HR-MYCN_A) (n = 33), High-risk MYCN not-amplified (NBL-HR-MYCN_NA) (n = 91), and Intermediate and Low-risk MYCN not-amplified (NBL-ILR) (n = 26), because these subgroups were shown to have distinct immune infiltration (Wei et al. 2018) (P. Zhang et al. 2017). Dimension-reduction using t-Distributed Stochastic Neighbor Embedding (tSNE) based on the whole transcriptome also showed separation of NBL subgroups (**Figure 1A**), therefore, we considered each subgroup as a separate cancer type in our analysis.

**Figure 1:**
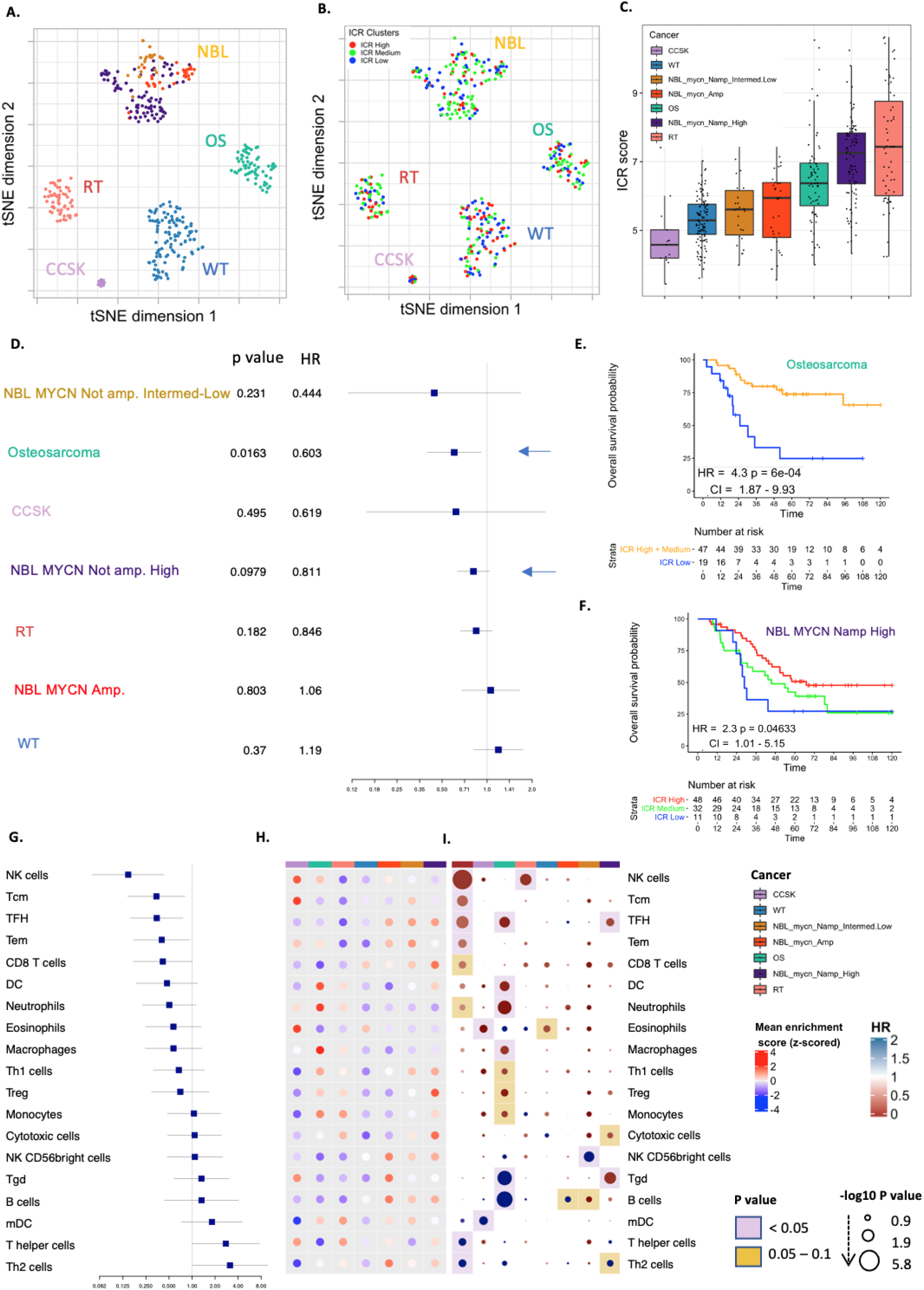
Prognostic role of immune signatures in pediatric tumors. (A) tSNE plot of filtered normalized expression values annotated by pediatric cancer types. (B) tSNE plot of filtered normalized expression values annotated by ICR clusters. (C) Boxplot of the ICR score across pediatric tumor types. (D) Forest plot of the association of continuous ICR scores with survival across tumors (E) Kaplan-Meier overall survival curve for ICR High + medium (orange) versus ICR low (blue) in Osteosarcoma. (F) Kaplan-Meier overall survival curve for ICR High versus ICR low in NBL-HR-MYCN_NA. (G) Forest plot of Hazards ratio of immune subpopulations across different pediatric tumors. (H) Heatmap of enrichment scores of immune cells signatures. (I) Hazards ratio heatmap of these signatures, the color of the circle representing the HR; HR below 1 is red and above 1 is blue, the radius size representing the −log10 p value; Larger size has higher −log10 p value and more significant association with survival, the color of the background corresponding to the p value; if pink; p value is less that 0.05, if yellow; p value between 0.05 and 0.1 and the white means p value above 0.1.

We used the expression of ICR genes to classify the overall immune orientation of cancer samples. The ICR genes exhibited high overall correlation with each other in most TARGET pediatric solid tumor cohorts, with a lower correlation in NBL-HR-MYCN_A and in WT (**Supplementary Figure 1**). Samples from each cancer type were separated into three ICR clusters (“ICR High”, “ICR Medium” and “ICR Low”). While dimension reduction of the expression data shows no clear segregation of samples by ICR clusters within each tumor type (**Figure 1B**), a clear difference in the distribution of ICR scores across cancer types was observed (**Figure 1C**). Lower ICR scores were found in WT and CCSK, while RT had the highest ICR scores. Significant differences in ICR scores were observed across NBL subgroups (p < 0.00001) for NBL-HR-MYCN_NA vs NBL-ILR and NBL-HR-MYCN_A respectively (**Supplementary Figure 2B**), reflecting large immune orientation differences between samples within NBL. Substantially lower ICR scores were observed in NBL-HR-MYCN_A and in NBL-ILR when compared to NBL-HR-MYCN_NA. This finding is consistent with previous reports of poor T-cell infiltration in NBL-HR-MYCN_A (Layer et al. 2017)(Wei et al. 2018) (P. Zhang et al. 2017) and higher T-cell infiltration in NBL-HR-MYCN_NA (Wei et al. 2018). Overall survival analysis of continuous ICR scores showed significant association of ICR scores with high survival rate in Osteosarcoma (p <0.016) (**Figure 1D**) and comparing the survival between the ICR clusters showed that in Osteosarcoma (OS), the ICR Low group had significantly lower overall survival (p < 0.001) (**Figure 1E**) and event-free survival estimates (p < 0.05) (**Supplementary Figure 2D**) compared to the other groups combined. The same pattern was found in NBL-HR-MYCN_NA where ICR High was associated with better overall survival compared to ICR Low (**Figure 1F**). This pattern was reversed in WT (**Supplementary Figure 2C**), as has been observed in adult kidney tumors (Roelands et al. 2020). No significant association with survival was found in rhabdoid tumor or clear cell sarcoma of kidney (Kaplan Meier curve for CCSK not showed because of the low number of samples n=13).

Since ICR was most prognostic in Osteosarcoma and high-risk MYCN-not-amplified NBL, we proceeded to examine which tumor intrinsic attributes correlate with immune infiltration, reflected by ICR score, in OS and NBL-HR-MYCN_NA. Tumor intrinsic pathways that correlated with ICR score in these tumors (**Supplementary Figure 3A**) are TNFR1, PI3K Akt mTOR signaling, Immunogenic cell death, Apoptosis, mTOR and others, while signatures inversely correlated with survival in both tumors include barrier genes, mismatch repair, proliferation, G2M checkpoints. Wnt beta catenin signaling showed very strong inverse correlation with ICR in Osteosarcoma but not in MYCN not amplified NBL (**Supplementary Figure 3A**).

We then examined the association of tumor intrinsic attributes with survival in these tumors, Wnt beta catenin pathway was significantly associated with a worse prognosis in OS (p < 0.05). We did not observe this same association in NBL-HR-MYCN_NA (**Supplementary Figure 3C**). In this neuroblastoma subgroup, several pathways were associated with worse prognosis, such as Myc targets, Glycolysis, mTORC1, DNA repair, Mismatch repair, E2F targets, G2M checkpoints and proliferation (**Supplementary Figure 3C**).

### Immune subpopulations show prognostic value in pediatric tumors

To explore the different immune characteristics of pediatric tumor types in more depth, we compared the enrichment of leucocyte subpopulations within and among cancer types (pan-cancer), using the gene expression signatures of previously published datasets (Bindea et al. 2013)(Angelova et al. 2015), as described in the methods section. Signatures such as NK cells, Tcm, TFH, Tem, CD8+ T-cells and neutrophils were significantly associated with better overall survival in the pan-cancer analysis, while T helper and Th2 cells were associated with worse prognosis (**Figure 1G**).

Compared with other cancer types, Osteosarcoma showed an immune active phenotype illustrated by increased mean enrichment of transcripts for dendritic cells (DC), macrophages, neutrophils, and mDC (**Figure 1H**). Enrichment scores for some leucocyte populations were associated with significantly improved prognosis as TFH, DC, neutrophils, macrophages, monocytes, Th1 and regulatory T cells (Treg) (**Figure 1I**), while B cell and gamma delta T-cell enrichments were associated with significantly worse survival in this cancer type (**Figure 1I**).

In Neuroblastoma, T cells, CD8+ T cells, Th17, NKT cells, Th1 cells, Treg cells, and DCs were significantly higher enriched in the high-risk MYCN-not-amplified group compared to MYCN amplified NBL (p < 0.05), TFH was high in the 3 subgroups and showed significant positive association with survival in the NBL-HR-MYCN_NA group. Gamma delta T cell (Tgd) enrichment also showed high association with survival in High-risk MYCN not-amplified group (p < 0.05). However, Th2 cells and NK CD56 bright cells were significantly higher enriched in the *MYCN* amplified group compared to NBL-HR-MYCN_NA group (p < 0.05) and in NBL-ILR, strong association of NK CD56 bright cells with worse prognosis was seen (p < 0.05).

In kidney tumors, WT and CCSK were characterized by low immune infiltration illustrated by a low ICR score, while RT showed the highest ICR score (**Figure 1B**). Decreased infiltration was associated with better survival in WT (**Supplementary Figure 1C**); this reverse association was previously observed in adult kidney cancer (Roelands et al. 2020). Overall low enrichment of immune subpopulations was observed in WT with no clear association with survival (**Figure 1H, 1I**).

In pan-cancer, expression patterns consistent with enrichment of several immune cells were associated with favorable prognosis, including NK cells, Tcm, TFH, Tem, CD8 T cells and Neutrophils, while the pattern was reversed in Th2 and T helper cells, consistent with similar observations in adult cancer. However, due to the small sample size of some cohorts, we could not identify consistent significant prognostic biomarkers in the leukocyte populations across all cancers.

### Immune subtypes show different immune characteristics in pediatric tumors

To further elucidate the impact of the cancer immune phenotypes in pediatric solid cancer, we expanded our analysis to a collection of previously published immune signatures. We performed ssGSEA on 105 immune signatures and clustered them to define modules of highly correlated immune signatures. We identified 5 main clusters of signatures (5 modules). Interestingly, in each of these modules we could identify one of the representative signatures presented in Thorsson et al (Thorsson et al. 2018) (INF-G, TGF-B, Macrophages, Lymphocytes, Wound healing) (**Figure 2A**). This finding demonstrates that modules of immune signatures in pediatric cancer showed a similar pattern of correlation as those identified in adult solid tumors, reflecting the robustness of these modules.

**Figure 2:**
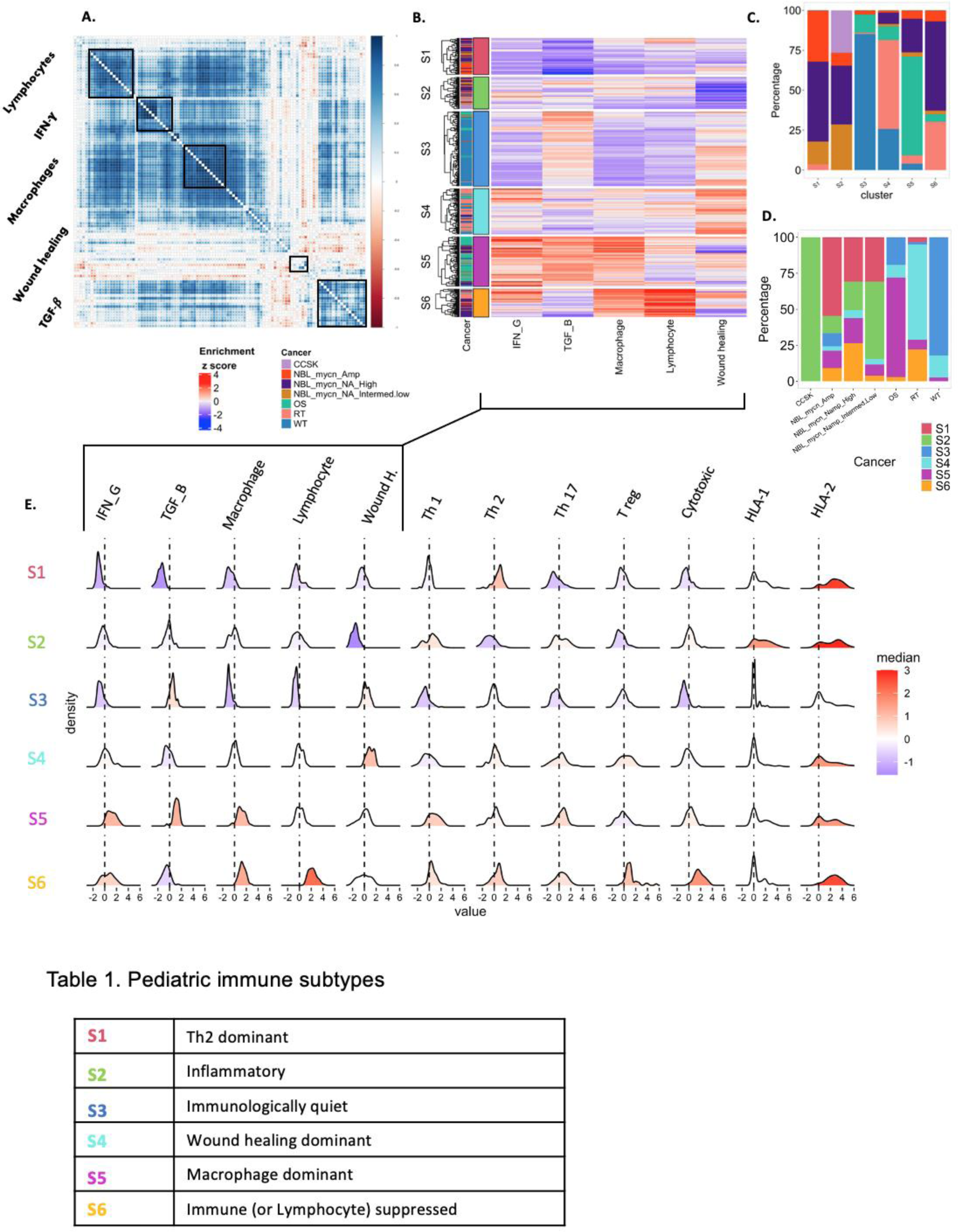
Immune subtypes of pediatric tumors. (A) Spearman’s correlation of 105 cancer immune signatures. Highly correlated signatures are clustered into 5 modules (black rectangles). (B) Heatmap showing the enrichment of immune signatures for each patient. Tumors are clustered into 6 subtypes based on the enrichment patterns. (C) Distribution of immune subtypes within TARGET pediatric tumors. Colors represent immune subtypes. (D) Distribution of cancer types within immune subtypes. Colors represent cancer types. (E) Distributions of signature scores within the six immune subtypes (rows), with dashed line indicating the median.

We then clustered the 408 patients based on the enrichment scores of these 5 representative immune gene signatures, into 6 immune subtypes (S1 to S6) with distinct immunologic orientation (**Figure 2B**). Each subtype includes patients from several tumor types (**Figure 2C**), and each tumor type consists of different immune subtypes (**Figure 2D**). We generated density plots of each of the five representative immune signatures and also of each of seven additional immune biomarkers that are known to reflect the immune orientation (**Figure 2E**). This allowed us to better interpret each immune subtype and label them based on their enrichment profiles (**Figure 2F**). S1 is referred to as Th2 dominant subtype as it has the highest Th2 and the lowest TGF-B, Macrophage, Lymphocyte and IFN-g signal. S2 was labeled the Inflammatory subtype since it exhibits the highest Th1-Th2 ratio, the highest HLA1 expression and lowest wound healing enrichment. Since high TGF-B stands out in S3, in addition to the low enrichment of Th1, Th17 we call this subtype Immunologically quiet. S4 or the Wound healing subtype is dominated by the Highest wound healing enrichment, shows high Th2 and Treg cells presence. The S5 subtype has increased TGF-B and IFN-G signatures, high Th1 and Th17 enrichment but seems to be immunologically impaired by high Macrophage presence and is thus referred to as Macrophage dominant. The last immune subtype, S6 or the Lymphocyte suppressed subtype, has enrichment of almost all properties of a high immune infiltration including counter regulatory signals from Th2, Treg, downregulated HLA1, and Macrophage presence. However, it is also characterized by high expression of immune checkpoints and exhaustion markers, so we call it immune (or lymphocyte) suppressed. (**Figure 2E**).

### Immune Cellular Fraction Estimates (CIBERSORTx)

We compared the immune cell fractions across the immune subtypes (**Supplementary Figure 5, Supplementary Figure 6**). High proportions of macrophages were observed in S5, with increased M2 macrophages proportions in S4, high proportions of mast cells observed in S3 and elevated proportions of lymphocytes were found in S2 that has the highest survival. In addition, it was very clear the high proportions of lymphocytes in S6 that characterized by the elevation of immune checkpoints and exhaustion markers which suppress the effect of T lymphocytes and make it in an exhaustion status (**Figure 4B**).

### Immune subtypes showed difference in overall survival

Cox proportional hazard models demonstrated significant violation of the model when adding the cancer type as a co-variate, so we stratified the model for the cancer type and performed Cox proportional hazard regression analysis which showed significant differences in overall survival between subtypes (**Figure 3A**). The best prognosis was observed for the Inflammatory subtype (S2) while the subtype with the worst survival was the Wound healing dominant subtype (S4). The lowest enrichment of Wound healing signature observed in S2, the subtype with the best survival among other immune subtypes (**Figure 3A**), suggesting an association of Wound healing signature expression with prognosis in pediatric tumors. In order to assess whether the difference in survival across immune subtypes is due to the tumor type distribution between clusters, we performed multivariate analysis using a Cox proportional hazards model including the cluster (immune subtype) and cancer type as co-variants, and we again found a significant difference in survival between S2 and S4 (p = 0.02), between S5 and S4 (p= 0.013), and between S6 and S4 (p = 0.0325), demonstrating the prognostic impact of this immune stratification (**Figure 3B**).

**Figure 3:**
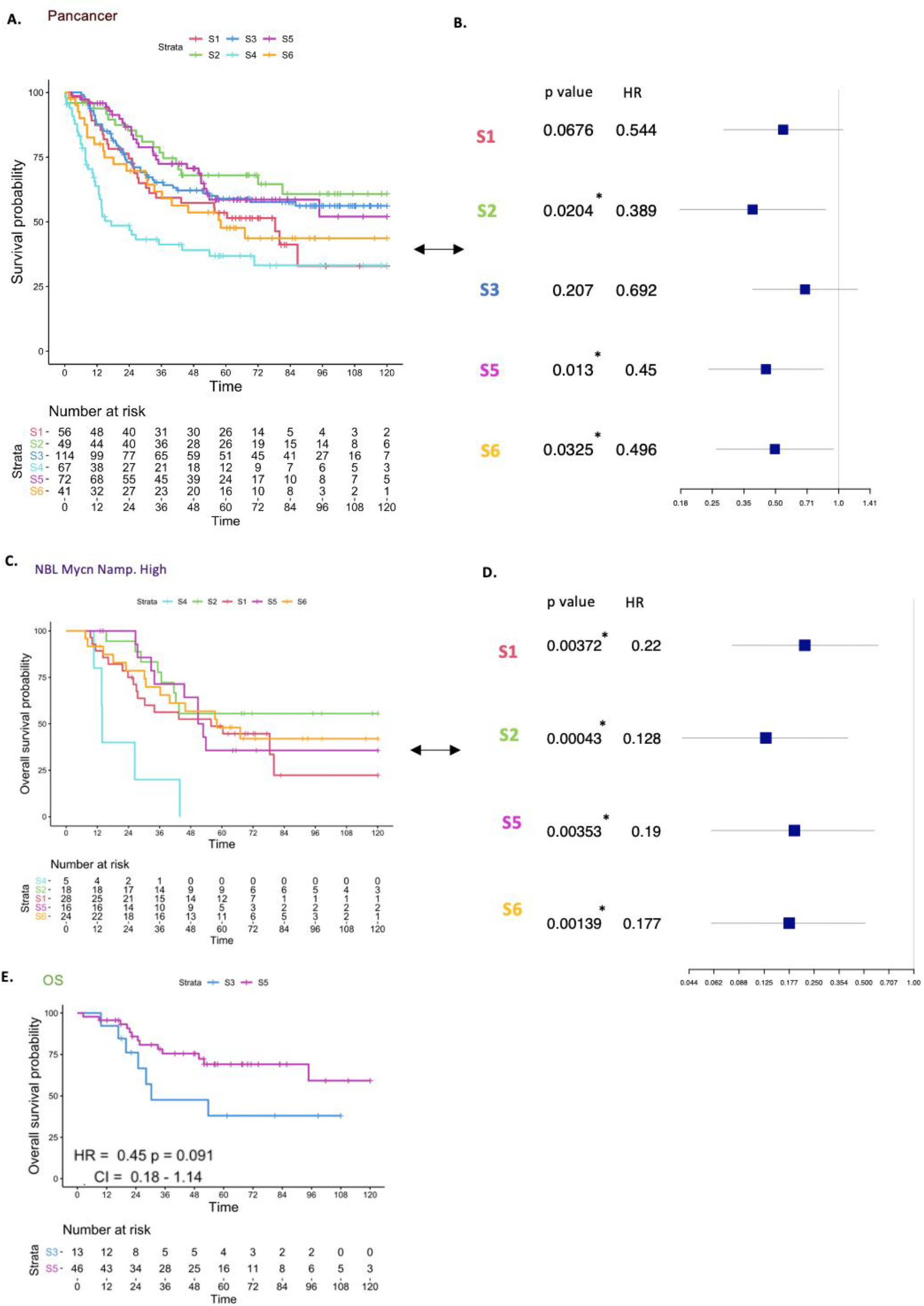
Overall survival across Immune subtypes. (A) Kaplan-Meier overall survival curve for immune subtypes. (B) Forest plot showing HRs (overall survival) of immune subtypes; S1, S2, S3, S5, S6 versus S4, and p value corrected for cancer types. (C) Kaplan-Meier overall survival curves for immune subtypes within tumor types for NBL-HR-MYCN_NA. (D) Forest plot showing HRs (overall survival) of immune subtypes within high-risk NBL-HR-MYCN_NA; S1, S2, S5, S6 versus S4. (E) Kaplan-Meier overall survival curves for immune subtypes within Osteosarcoma.

In order to test the prognostic value of our immune stratification within each tumor type, we compared overall survival between immune subtypes within each tumor (**Figures 3C-E, Supplementary Figure 4**). Interestingly, for NBL-HR-MYCN_NA tumors, we found significant differences between all the immune subtypes versus S4 (p < 0.05) which indicates the presence of a subgroups with distinct immunological features within the NBL-HR-MYCN_NA cohort. The same survival pattern was observed for S4 in both Wilms and Rhabdoid tumors, and a clear difference in survival between the S3 and S5 subtypes was seen in Osteosarcoma (p= 0.09) (**Figure 3E**). For CCSK the number of samples was too small and therefore the K-M curve was not plotted. These observations highlight the immune heterogeneity within tumors and the importance of understanding the immunological features of pediatric tumors and their subgroups in order to raise the therapeutic effect.

NBL is heterogenous tumor and different clinical parameters contribute to the survival of NBL (Wei et al. 2018), (Z. Liu et al. 2020), as previously mentioned, significant difference across immune subtypes within the NBL-HR-MYCN_NA was observed (**Figure 3C, 3D**), we performed multivariate analysis to correct for the contribution of other clinical parameters in the survival of NBL, significant difference in survival was found between S2 (p= 0.0319) and S6 (p= 0.0452) compared to S4.

### Immune checkpoints expression in immune subtypes

To understand the prognostic role of immune checkpoints in pediatric tumors, we performed survival analysis for checkpoint expression pan-cancer and across our immune subtypes. The *CD276* gene was significantly associated with survival in pan-cancer analysis (**Figure 4A**). We noticed that immune checkpoints that are strongly associated with better prognosis as *CD276, KIRD3DL1, VTCN1, C10orf54 (VISTA)*, had a low enriched in S4 (**Figure 4B**), while those associated with worse prognosis, the immune checkpoints as *LAG3, CD70*, TNFSF4, IDO1, *KIRD3DX1, CD28 and TNFSF9*, were highly expressed in S4. To deeper understand the prognostic effect of the immune checkpoints expression within each immune subtype we generated a HR heatmap annotated by immune subtypes (**Figure 4C**). In S4 a unique pattern of association with worse prognosis was found with *C10orf54 (VISTA)* and *CD86* (p 0.05 – 0.1). While *TNFRSF9* was associated with worse survival in S4 and S3. In pan-cancer significant association of *CD70* and *LAG3* expression with poor survival was observed (p < 0.05). Some immune checkpoints show reverse pattern of survival with different subtypes as *C10orf54* that associate with reverse favorable prognosis in S2, *TNFRSF14* across S1 and S3 and *TNFRSF4* across S2 and S6. These findings reflect the variation in prognosis of immune checkpoints expression within different tumor types.

**Figure 4:**
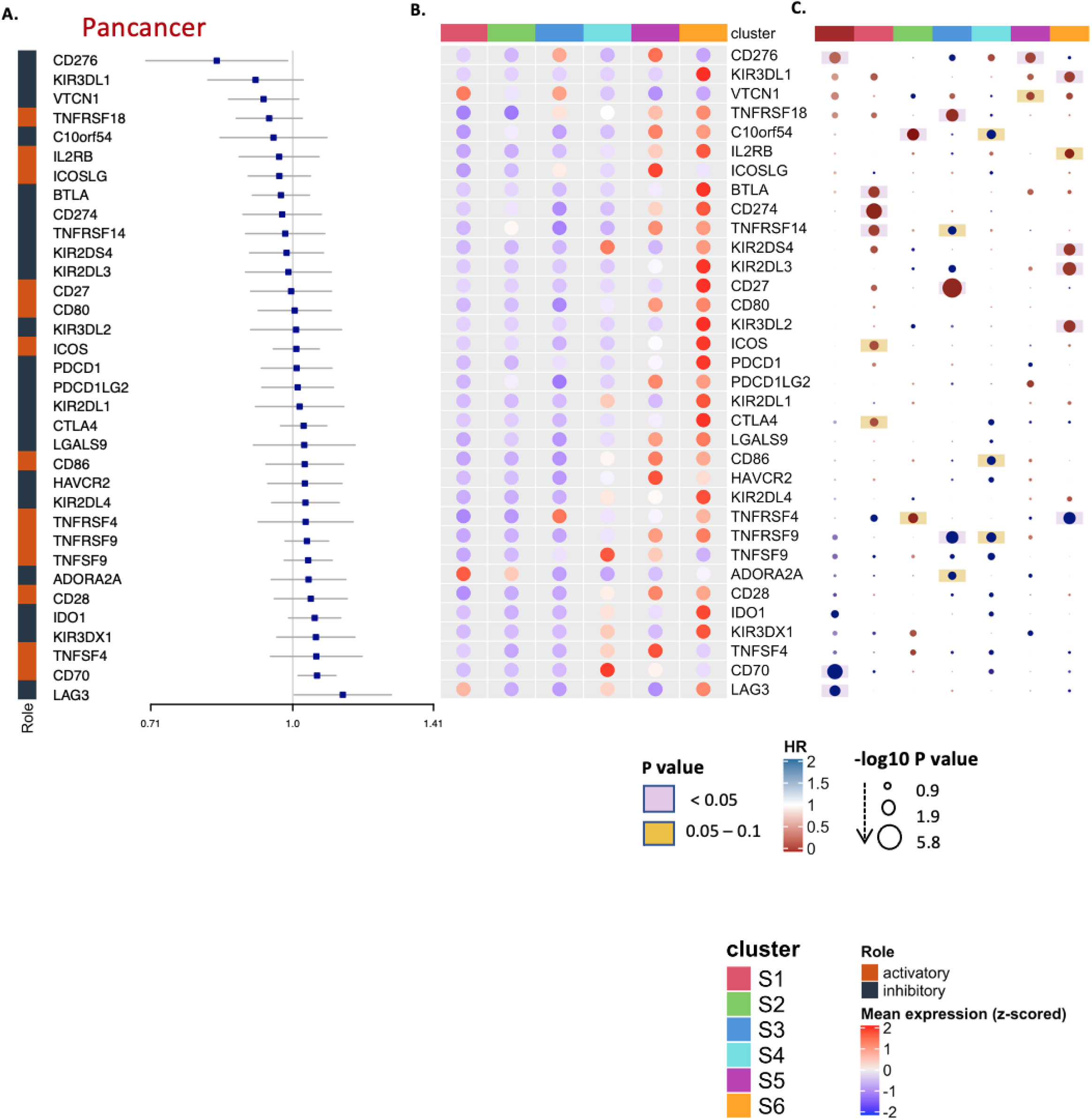
Immune checkpoints expression across Immune subtypes. (A) Pan-cancer forest plot showing HRs (overall survival) and p-values of immune checkpoint expression. (B) Heatmap of log2 transformed expression of immune checkpoints aggregated by the median for each immune subtype, split into activator and inhibitor and annotated by immune subtypes. (C) Hazard ratio heatmap of these immune checkpoints, the color of the circle representing the HR; HR below 1 is red and above 1 is blue, the radius size representing the −log10 p value; Larger size has higher −log10 p value and more significant association with survival, the color of the background corresponding to the p value; if pink; p value is less than 0.05, if yellow; p value between 0.05 and 0.1 and the white means p value above 0.1. (D) Violin plots illustrating the difference in expression of selected immune checkpoints across S2 and S4 immune subtypes.

High expression of immune checkpoints was observed in the Leukocyte dominant subtype (S6) compared to other immune subtypes (**Figure 4B**). This could be explained by the exhaustion status seen in this immune subtype which displays the highest enrichment of lymphocytes (**Figure 2E**).

### Selected oncogenic pathways and the immune phenotypes

We also examined tumor intrinsic differences between immune subtypes, by investigating the association between overall survival and the expression of tumor intrinsic pathways in pan-cancer and across the immune subtypes (**Figure 5A, 5C**), and comparing the enrichment of tumor intrinsic pathways between the 6 immune subtypes (**Figure 5B**). A wide variety of pathways were differentially enriched between immune subtypes. Myc targets, DNA repair and oxidative phosphorylation showed a uniquely high enrichment in S4 compared to other groups. However, Wnt beta catenin and TGF-B showed a similar enrichment pattern among immune subtypes with increased enrichment in S3 and S5 (**Figure 5B**). Interestingly, most of the pathways show a mirrored expression level between S2 and S4; for example, S4 was significantly higher in the enrichment of TGF-B and Barrier genes compared to S2, while p38 MAPK Signaling, ErbB2 ErbB3 Signaling, NOS1 Signature and SHC1/pSTAT3 Signatures were significantly highly enriched in S4 vs. S2 (p < 0.05). Within the immune subtypes, significantly high association of some oncogenic pathways with worse prognosis seen exclusively in S4 as mTORC1, Myc targets, NOS1, ERK5, PI3K AKT. (**Figure 5C**).

**Figure 5:**
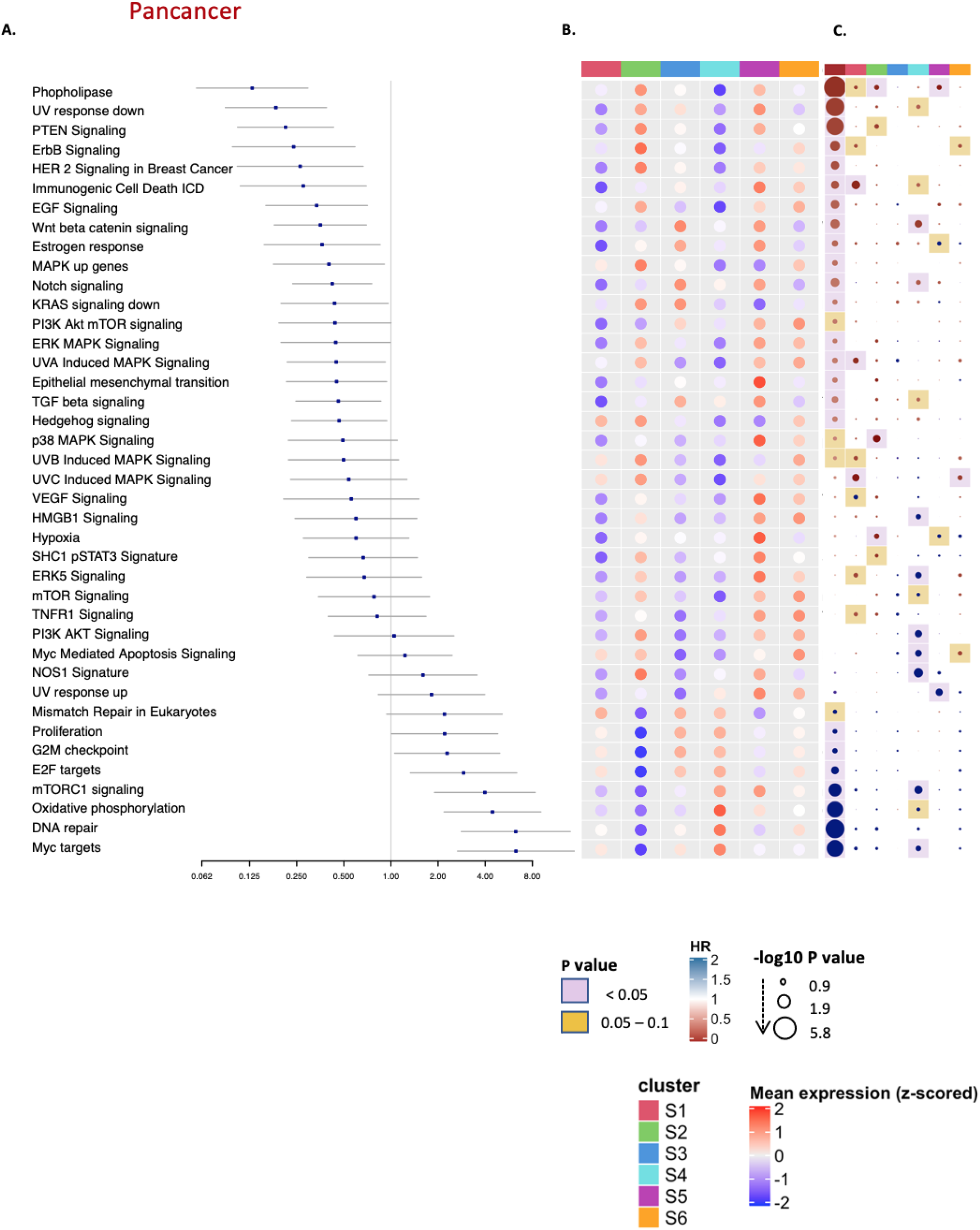
Intrinsic selected oncogenic pathways across immune subtypes. (A) Pan-cancer forest plot showing HRs (overall survival) and p-values of selected oncogenic pathway enrichment scores. (B) Heatmap of the enrichment scores of selected oncogenic pathways, blue colors corresponding to lower enrichment and the red for the high enrichment scores, annotated by the 6 immune subtypes and pan-cancer. (C) Hazards ratio heatmap of these pathway enrichment scores, the color of the circle representing the HR; HR below 1 is red and above 1 is blue, the radius size representing the −log10 p value; Larger size has higher −log10 p value and more significant association with survival, the color of the background corresponding to the p value; if pink; p value is less than 0.05, if yellow; p value between 0.05 and 0.1 and the white means p value above 0.1.

## Discussion

Understanding the relationship between tumor cells and the host’s immune system has proven to be instrumental in advancing the treatment of adult cancers. It is self-evident that the role of the immune system in pediatric cancer can no longer be ignored. In this study we set out to transpose the knowledge gained in the transcriptomic analysis of the TARGET data. It is fundamental to gain a deeper understanding of the prognostic connotations of the immune phenotypes present in pediatric solid tumors, in order to evolve more effective therapeutic strategies and stratification systems. Here we use a range of transcriptional signatures to investigate the composition, proportion, activation, and functional orientation of immune infiltrates within the TME of pediatric tumors and the interaction of these immune infiltrates with the oncogenic pathways of these tumors.

We performed a pan-cancer immunogenomic analysis and found that the disposition of an immune active phenotype characterized by T helper 1 immune infiltration was associated with favorable prognosis in Osteosarcoma and NBL-HR-MYCN_NA. The overall immune phenotype was determined by a gene signature we have used in the context of adult cancer on several occasion that has been shown to have strong association with prognosis and response to immunotherapy and that captures well the presence of active immune mediated tumor rejection. (Hendrickx et al. 2017), (Roelands et al. 2020) The Immunologic Constant of Rejection showed a prognostic role in pediatric Osteosarcoma in which high immune infiltration, present in ICR High clustered samples, where it was associated with better prognosis. This finding is in agreement with results obtained from Zhang et al. (C. Zhang et al. 2020) in which tumors with high immune score that estimate immune cell infiltration by analyzing a certain gene expression signature of TIICs, were associated with higher survival rate than tumors with low immune score in Osteosarcoma.

WNT/β-catenin signaling also showed a prognostic role in OS, we found an inverse correlation of Wnt beta catenin pathway enrichment with immune infiltration (ICR score) in OS, WNT/β-catenin signaling in melanoma is an essential tumor intrinsic oncogenic pathway associated with an immune inactive phenotype (Fu et al. 2015, 10), (Spranger and Gajewski 2015). A recent study by Luke etal. (Luke et al. 2019) confirmed this finding in several adult tumors obtained from TCGA, they demonstrated the inverse correlation between β-catenin protein level and T-cell-inflamed gene expression. In addition to the association with low immune infiltration, WNT/β-catenin signaling enrichment was significantly associated with significant worse outcomes in Osteosarcoma. A meta-analysis performed by Xie et al. confirmed the association of over-expression of β-catenin with metastasis in Osteosarcoma. (Xie et al. 2020) Several clinical trials are ongoing the combination of immunotherapy with WNT/β-catenin signaling inhibitors for the treatment of various tumors(Y. Zhang and Wang 2020).

In High-risk Neuroblastoma, despite the introduction of anti–GD2 antibody-based immunotherapy, the cure rates remain low, and there is also significant morbidity in survivors (Kudva and Modak 2019), (Yust-Katz, Khagi, and Gilbert 2020). As seen in OS, we demonstrated that NBL-HR-MYCN_NA was associated with significantly higher immune infiltration, represented by ICR score compared to NBL-HR-MYCN_NA tumors, confirming what published in 2018 by Jun S. Wei (Wei et al. 2018) who found that *MYCN* not-amplified (MYCN-NA) tumors had significant higher cytotoxic TIL signatures compared to *MYCN* amplified (MYCN-A) tumors, or previously in 2017 by Zhang et al. (P. Zhang et al. 2017) who demonstrated the association of *MYCN* amplification with repressed cellular immunity, in addition to Layer et al (Layer et al. 2017) that demonstrated the association of T cell poor microenvironment in primary metastatic neuroblastomas that exhibit lower interferon pathway activity and chemokine expression with genomic amplification of the *MYCN (N-Myc)* proto-oncogene.

A strong association between the presence of T-cell-infiltration and/or a T-cell-inflamed gene expression signatures, and response to immune-checkpoint blockade has been previously observed (Chen et al. 2016), (Tumeh et al. 2014), (Hellmann et al. 2019), (Hamid et al. 2011). Increased response to immune checkpoints therapy after T cell activation/functioning was extensively described and explained by the production of *IFN* by tumor-infiltrating CD8 (Bald et al. 2014), (Taube et al. 2014), (Grywalska et al. 2018), that in turn induce the expression of PD-L1 in tumor cells, which is a well-studied predictive biomarker for response to immune checkpoint inhibitors (Doroshow et al. 2021).

In contrast to what was observed in OS, immune infiltration showed an association with worse prognosis in Wilms tumor. Interestingly, we previously reported clear cell renal cell carcinoma (ccRCC)(Roelands et al. 2020). This reverse association was also observed in a number of other adult tumors like Uveal Melanoma (UVM) and Low-Grade Glioma (LGG), in which a significant reverse association was observed between the ICR High patients and survival. The Tumor types in which this reverse association was observed are referred to as ICR Disabled tumor types and they are typified by low mean of ICR score, ICR high association with worse survival, low neoantigen load, down regulation of pathways involved in proliferation and low enrichment of *TGF-B* signature. (Roelands et al. 2020).

Since we established that the immune infiltrate does impact the pediatric cancer prognosis we deepened our analysis and demonstrated that solid pediatric cancer samples can be meaningfully immune subtyped, comparable to what was achieved in adult cancer (Thorsson et al. 2018). In the TARGET dataset, the same 5 representative signatures (INF-G, TGF-B, Macrophages, Lymphocytes and Wound healing) identified in Thorsson et al, were found to also be at the cores of the 5 main co-clustering modules of immune signatures (Thorsson et al. 2018). When we then clustered the TARGET samples based on the enrichment scores of these 5 representative signatures, although the low number of tumor samples that we have, we obtained 6 robust immune subtypes. These displayed distinct immune characteristics and significant difference in overall survival rate even after adjusting for cancer type. Some of these immune subtypes share similar immune features with adult tumors, like the ‘Immunologically quiet’ subtype that is characterized by overall low enrichment T cells and IFN-G signature, or the ‘inflammatory subtype’, that is characterized by high Th1 enrichment and is associated with the best survival across all immune subtypes in pediatric and adult tumors.

High enrichment of wound healing signatures was associated with worse outcomes in adult tumors (J. Liu et al. 2021), (Thorsson et al. 2018), we observed the same in pediatric tumors in which the wound healing dominant subtype (S4) showed the worst survival across the pediatric immune subtypes. Wound healing enrichment is accompanied by elevated expression of angiogenic genes, and high enrichment of proliferation signature. S4 also showed high proportions of Macrophages shown by the CIBERSORTx (**Supplementary Figure 6**), especially macrophages M2 (**Supplementary Figure 5**).

Different clinical parameters contribute to the survival in Neuroblastoma (Brodeur and Nakagawara 1992), (Campbell et al. 2020). In the TARGET dataset, the risk classification followed the Children Oncology Group (COG) that includes the age, histology, ploidy, *MYCN* status, and MKI as parameters for the stratification. (“Https://Www.Cancer.Org/Cancer/Neuroblastoma/Detection-Diagnosis-Staging/Risk-Groups.Html,” n.d.), (Sokol and Desai 2019)

Previous studies discussed the significant difference in immune characteristics across neuroblastoma subgroups and its impact on survival, Wei et al (Wei et al. 2018), showed that a cytolytic immune signature was associated with a favorable prognosis in NBL-HR-MYCN_NA and a high *MYC* expression signature was associated with worse prognosis in the same group. Similar data was obtained by Valentijn et al (Valentijn et al. 2012), Lee et al (J. W. Lee et al. 2018), who demonstrated the association of *MYCN* amplified NBL with relapse and progression. This analysis agreed with the previous findings, the prognostic value of immune phenotyping based on ICR and the 5 representative signatures showed prognostic value in the NBL-HR-MYCN_NA only. With the best prognosis in S2 that is characterized by high enrichment of inflammatory signatures and the lowest in S4 that is dominated by Wound healing, a similar survival pattern as observed pan-cancer.

Recent studies described the immunosuppressive checkpoint molecules that downregulate immune cell function in the aftermath of infection. These immune checkpoints can contribute to immune evasion and tumor escape when found on malignant cells or in the tumor microenvironment. Targeting immune checkpoints particularly Programmed cell Death protein 1 (*PD-1/CD279*), Programmed Death Ligand 1 (*PD-L1/CD274*) and Cytotoxic T-lymphocyte antigen-4 (*CTLA-4/CD28*) has made a breakthrough in treating advanced adult malignancies. However, there is limited investigations performed on immune checkpoint molecules and inhibitors in solid pediatric tumors (Kabir et al. 2018).

It was previously demonstrated that *IDO1* has an anti-inflammatory effect that counters the effect of activated T-cells in inflammatory bowel disease (Wolf 2004) and expression of *IDO1* was associated with recruitment and activation of myeloid-derived suppressor cells through a Treg-dependent mechanism that contributed to aggressive tumor growth and poor response to T-cell targeted therapy (Holmgaard et al. 2015), (Uyttenhove et al. 2003). Lymphocyte Activation Gene-3 (*LAG3*) has shown an immunomodulatory role through suppressing T-cell activation and cytokine secretion, and as a result, it maintains immune homeostasis. It was also found to drive a differential lymphocyte inhibitory effect and showed a synergetic effect with *PD-1* to suppress immune responses in cancer. Targeting *LAG-3* as immunotherapy is in clinical trial, and in combination immunotherapy, anti-LAG-3 and anti-PD-1 has demonstrated powerful efficacy in overcoming *PD-1* resistance (Gestermann et al. 2020). In our analysis, we observed high expression of *IDO1* and *LAG3* in the wound healing immune subtype (S4) which may both explain the poorer prognosis associated with this subtype and also and should be the first in line to be further explored in the pediatric setting.

Other immune checkpoints of note are Cluster of Differentiation 70 (*CD70*) and TNF Superfamily Member 9 (*TNFSF9*), they are co-stimulatory immune checkpoints and are members of the tumor necrosis factor (*TNF*) family (O’Neill et al. 2017), (Fröhlich et al. 2020). We found that they are highly expressed in S4 compared to other immune subtypes, these checkpoints are associated with NK cells innate immunity (Takeda et al. 2000). *CD70* has been shown to correlate with worse relapse-free survival rates in breast cancer patients, that could be due to the unfavorable negative feedback function to downregulate inflammatory T cell responses obtained by T cell-derived *CD70* (O’Neill et al. 2017). Significant differences in the expression of immune checkpoints across the immune subtypes, and the different association of immune checkpoints with survival highlight the value of stratifying pediatric solid tumors into different immune phenotypes and allow us to understand why the response to immune treatments varies across patients even within the same tumor type. Further investigation will be needed to address the importance of these findings towards the possible implementation of check point inhibition studies in pediatric cancer.

Next to the expression of checkpoints we noted a significantly higher enrichment of DNA repair signature pathways compared to other immune subtypes in S4 (**Figure 5B**). DNA mismatch and DNA repair mechanisms in addition to microsatellite instability are well known biomarkers for response to immune checkpoints (Teo et al. 2018), (Jiang et al. 2021), (Le et al. 2017). Defects in DNA damage response can radically influence the equilibrium between immune surveillance and tumor progression. On one hand, erroneous DNA damage repair results in the buildup of mutations or chromosomal rearrangements, that promote genomic instability, oncogene activation and initiation of tumorigenesis. On the other hand, it can trigger anti-tumor immune response through neo-antigens production that are later presented on the cell surface in the context of major histocompatibility complex class I and detected by the immune cells and the increase of cytosolic DNA that in turn stimulates the innate immune response (J. Zhang, Shih, and Lin 2020), (Cerniglia et al. 2019).

In addition, DNA repair pathways defects in different cancers also provide therapeutic opportunities to kill cancer cells without affecting normal cells as with the use of poly(ADP ribose) polymerase (PARP) inhibitors for the treatment of homology-dependent recombination defective cancers (Farmer et al. 2005)(Bryant et al. 2005)(Rouleau et al. 2010)

In this paper, we demonstrated that extracranial solid pediatric tumor can be classified according to their immune disposition, unveiling unexpected similarity with adults’ tumors. Immunological parameters can be further explored to refine diagnostic and prognostic biomarkers and to identify potential immune responsive tumors. This is the first pan-cancer immunogenomic analysis in children.

### Limitations of the study

In our study we used 408 tumors for stratifying the cohort into immune subtypes, which is considered low in comparison to TCGA, this issue was prominent in some small cohorts that affects the significance of the results, however; In spite of this small cohort we could see many significant differences across the phenotypes that we have, so validation of the analysis on different cohorts is important. In addition to that, the absence of mutation data that obscure the genes expression of the tumor or the host immune cells.

## Abbreviations

CCSK: Clear Cell Sarcoma of Kidney
COG: Children’s Oncology Group
ES: Enrichment score
ICR: immunologic constant of rejection
mDC: Macrophage-derived chemokine
MKI: Mitosis-Karyorrhexis Index
NBL: Neuroblastoma
NBL-HR-MYCN_A: High risk MYCN amplified Neuroblastoma
NBL-HR-MYCN_NA: High-risk MYCN not-amplified Neuroblastoma
NBL-ILR: Intermediate and low risk MYCN not amplified Neuroblastoma
OS: Osteosarcoma
RT: Rhabdoid tumor
WT: Wilms tumor

## Acknowledgement

“The results published here are in whole or part based upon data generated by the Therapeutically Applicable Research to Generate Effective Treatments (TARGET) initiative, phs000218. The data used for this analysis are available at https://portal.gdc.cancer.gov/projects.”

## Author contributions

S.S. Collected the public data, performed the bioinformatc analysis and drafted the manuscript. J.R., W.M., and E.A. participated in the interpretation of the results and edited the manuscript. D.B. and B.M participated in the conceptualisation and interpretation of the analysis and critically reviewed the manuscript. W.H. Coordinated the conceptualisation, analysis, interprestation and editing process.

## Ethics approval and consent to participate

Not applicable.

## Data availability statement

All scripts will be deposited in a Github release on Zenodo.

## Supplementary Figures

**Supplementary Figure 1:** Correlation of ICR genes in pediatric tumors. (A) Pearson correlation heatmap of ICR genes in different pediatric tumors, immune regulator genes were colored in blue and immune active genes in red, positive correlation between genes represented by blue and negative correlation represented by red.

**Supplementary Figure 2:** The Immunologic constant of rejection. (A) Pan-cancer heatmap of ICR gene expression annotated by cancer types, and per-cancer clustering ICR High, medium and low. (B) Boxplot showing the distribution of ICR scores in NBL-HR-MYCN_A, NBL-HR-MYCN_NA, and NBL-ILR (the p value was calculated by two-tailed t-test). (C) Kaplan-Meier of overall survival for ICR High versus ICR low in Wilms tumor. (D) Kaplan-Meier event free survival curve for ICR High + medium (orange) versus ICR low (blue) in Osteosarcoma. (E) Kaplan-Meier of overall survival for ICR High versus ICR low in Rhabdoid tumor.

**Supplementary Figure 3:** Intrinsic oncogenic pathways and immune infiltration. (A) Heatmap of pearson correlation between enrichment score (ES) of oncogenic pathways and ICR in pan-cancer, Osteosarcoma and NBL-HR-MYCN_NA. (B) Forest plot of HR of oncogenic pathways enrichment in Osteosarcoma (C) Forest plot of HR of oncogenic pathways enrichment in NBL-HR-MYCN_NA.

**Supplementary Figure 4:** Overall survival across Immune subtypes. (A) Kaplan-Meier overall survival curve for immune subtypes within Wilms tumor. (B) Kaplan-Meier overall survival curve for immune subtypes within Rhabdoid tumor. (C) Kaplan-Meier overall survival curve for immune subtypes within NBL-HR-MYCN_A tumors. (D) Forest plot showing HRs (overall survival) of immune subtypes within MYCN amplified NBL tumors; S1, S2, S3, S5 versus S6 (E) Kaplan-Meier overall survival curve for immune subtypes within NBL-ILR tumors.

**Supplementary Figure 5:** CIBERSORTx immune cells proportions across Immune subtypes. (A) Barplot of the median of proportions of CIBERSORTx immune cells in the 6 immune subtypes. (B) Boxplots of means of CIBERSORTx immune cells across the immune subtypes.

**Supplementary Figure 6:** CIBERSORTx immune cells proportions (Aggregate) across Immune subtypes. (A) Barplot of the median of proportions of aggregate CIBERSORTx immune cells in the 6 immune subtypes.

## References

Adamson, Peter C. 2015. “Improving the Outcome for Children with Cancer: Development of Targeted New Agents.”CA: A Cancer Journal for Clinicians 65 (3): 212–20. https://doi.org/10.3322/caac.21273.

Angelova, Mihaela, Pornpimol Charoentong, Hubert Hackl, Maria L Fischer, Rene Snajder, Anne M Krogsdam, Maximilian J Waldner, et al. 2015. “Characterization of the Immunophenotypes and Antigenomes of Colorectal Cancers Reveals Distinct Tumor Escape Mechanisms and Novel Targets for Immunotherapy.”Genome Biology 16 (1). https://doi.org/10.1186/s13059-015-0620-6.

Bald, Tobias, Jennifer Landsberg, Dorys Lopez-Ramos, Marcel Renn, Nicole Glodde, Philipp Jansen, Evelyn Gaffal, et al. 2014. “Immune Cell–Poor Melanomas Benefit from PD-1 Blockade after Targeted Type I IFN Activation.”Cancer Discovery 4 (6): 674–87. https://doi.org/10.1158/2159-8290.CD-13-0458.

Bertucci, François, Pascal Finetti, Ines Simeone, Wouter Hendrickx, Ena Wang, Francesco M. Marincola, Patrice Viens, et al. 2018. “The Immunologic Constant of Rejection Classification Refines the Prognostic Value of Conventional Prognostic Signatures in Breast Cancer.”British Journal of Cancer, October. https://doi.org/10.1038/s41416-018-0309-1.

Bindea, Gabriela, Bernhard Mlecnik, Marie Tosolini, Amos Kirilovsky, Maximilian Waldner, Anna C. Obenauf, Helen Angell, et al. 2013. “Spatiotemporal Dynamics of Intratumoral Immune Cells Reveal the Immune Landscape in Human Cancer.”Immunity 39 (4): 782–95. https://doi.org/10.1016/j.immuni.2013.10.003.

Brodeur, Garrett M., and Akira Nakagawara. 1992. “Molecular Basis of Clinical Heterogeneity in Neuroblastoma:”Journal of Pediatric Hematology/Oncology 14 (2): 111–16. https://doi.org/10.1097/00043426-199205000-00004.

Bryant, Helen E., Niklas Schultz, Huw D. Thomas, Kayan M. Parker, Dan Flower, Elena Lopez, Suzanne Kyle, Mark Meuth, Nicola J. Curtin, and Thomas Helleday. 2005. “Specific Killing of BRCA2-Deficient Tumours with Inhibitors of Poly(ADP-Ribose) Polymerase.”Nature 434 (7035): 913–17. https://doi.org/10.1038/nature03443.

Buchbinder, Elizabeth I., and Anupam Desai. 2016. “CTLA-4 and PD-1 Pathways: Similarities, Differences, and Implications of Their Inhibition.”American Journal of Clinical Oncology 39 (1): 98–106. https://doi.org/10.1097/COC.0000000000000239.

Burger, Jan A., Alessandra Tedeschi, Paul M. Barr, Tadeusz Robak, Carolyn Owen, Paolo Ghia, Osnat Bairey, et al. 2015. “Ibrutinib as Initial Therapy for Patients with Chronic Lymphocytic Leukemia.”New England Journal of Medicine 373 (25): 2425–37. https://doi.org/10.1056/NEJMoa1509388.

Campbell, Kevin, Arlene Naranjo, Emily Hibbitts, Julie M. Gastier-Foster, Rochelle Bagatell, Meredith S. Irwin, Hiroyuki Shimada, Michael Hogarty, Julie R. Park, and Steven G. DuBois. 2020. “Association of Heterogeneous MYCN Amplification with Clinical Features, Biological Characteristics and Outcomes in Neuroblastoma: A Report from the Children’s Oncology Group.”European Journal of Cancer 133 (July): 112–19. https://doi.org/10.1016/j.ejca.2020.04.007.

Cerniglia, Michael, Joanne Xiu, Axel Grothey, Michael J. Pishvaian, Jimmy J. Hwang, John Marshall, Ari M. Vanderwalde, et al. 2019. “Association of DNA Damage Response and Repair Genes (DDR) Mutations and Microsatellite Instability (MSI), PD-L1 Expression, Tumor Mutational Burden (TMB) in Gastroesophageal Cancers.”Journal of Clinical Oncology 37 (4_suppl): 60–60. https://doi.org/10.1200/JCO.2019.37.4_suppl.60.

Chen, Pei-Ling, Whijae Roh, Alexandre Reuben, Zachary A. Cooper, Christine N. Spencer, Peter A. Prieto, John P. Miller, et al. 2016. “Analysis of Immune Signatures in Longitudinal Tumor Samples Yields Insight into Biomarkers of Response and Mechanisms of Resistance to Immune Checkpoint Blockade.”Cancer Discovery 6 (8): 827–37. https://doi.org/10.1158/2159-8290.CD-15-1545.

Colaprico, Antonio, Tiago C. Silva, Catharina Olsen, Luciano Garofano, Claudia Cava, Davide Garolini, Thais S. Sabedot, et al. 2016. “TCGAbiolinks: An R/Bioconductor Package for Integrative Analysis of TCGA Data.”Nucleic Acids Research 44 (8): e71. https://doi.org/10.1093/nar/gkv1507.

Crist, W, E A Gehan, A H Ragab, P S Dickman, S S Donaldson, C Fryer, D Hammond, D M Hays, J Herrmann, and R Heyn. 1995. “The Third Intergroup Rhabdomyosarcoma Study.”Journal of Clinical Oncology 13 (3): 610–30. https://doi.org/10.1200/JCO.1995.13.3.610.

Dome, Jeffrey S., Elizabeth J. Perlman, and Norbert Graf. 2014. “Risk Stratification for Wilms Tumor: Current Approach and Future Directions.”American Society of Clinical Oncology Educational Book, no. 34 (May): 215–23. https://doi.org/10.14694/EdBook_AM.2014.34.215.

Doroshow, Deborah Blythe, b Sheena Bhalla, Mary Beth Beasley, Lynette M. Sholl, Keith M. Kerr, Sacha Gnjatic, Ignacio I. Wistuba, David L. Rimm, Ming Sound Tsao, and Fred R. Hirsch. 2021. “PD-L1 as a Biomarker of Response to Immune-Checkpoint Inhibitors.”Nature Reviews Clinical Oncology, February. https://doi.org/10.1038/s41571-021-00473-5.

Farmer, Hannah, Nuala McCabe, Christopher J. Lord, Andrew N. J. Tutt, Damian A. Johnson, Tobias B. Richardson, Manuela Santarosa, et al. 2005. “Targeting the DNA Repair Defect in BRCA Mutant Cells as a Therapeutic Strategy.”Nature 434 (7035): 917–21. https://doi.org/10.1038/nature03445.

Fröhlich, Anne, Sophia Loick, Emma Grace Bawden, Simon Fietz, Jörn Dietrich, Eric Diekmann, Gonzalo Saavedra, et al. 2020. “Comprehensive Analysis of Tumor Necrosis Factor Receptor TNFRSF9 (4-1BB) DNA Methylation with Regard to Molecular and Clinicopathological Features, Immune Infiltrates, and Response Prediction to Immunotherapy in Melanoma.”EBioMedicine 52 (February): 102647. https://doi.org/10.1016/j.ebiom.2020.102647.

Fu, Chunmei, Xinjun Liang, Weiguo Cui, Julia L. Ober-Blöbaum, Joseph Vazzana, Protul A. Shrikant, Kelvin P. Lee, Björn E. Clausen, Ira Mellman, and Aimin Jiang. 2015. “β-Catenin in Dendritic Cells Exerts Opposite Functions in Cross-Priming and Maintenance of CD8 + T Cells through Regulation of IL-10.”Proceedings of the National Academy of Sciences 112 (9): 2823–28. https://doi.org/10.1073/pnas.1414167112.

Galon, Jérôme, Helen K. Angell, Davide Bedognetti, and Francesco M. Marincola. 2013. “The Continuum of Cancer Immunosurveillance: Prognostic, Predictive, and Mechanistic Signatures.”Immunity 39 (1): 11–26. https://doi.org/10.1016/j.immuni.2013.07.008.

Gelband, Hellen, Prabhat Jha, Rengaswamy Sankaranarayanan, and Susan Horton, eds. 2015. Disease Control Priorities, Third Edition (Volume 3): Cancer. The World Bank. https://doi.org/10.1596/978-1-4648-0349-9.

Gestermann, Nicolas, Damien Saugy, Christophe Martignier, Laure Tillé, Silvia A. Fuertes Marraco, Markus Zettl, Iñigo Tirapu, Daniel E. Speiser, and Grégory Verdeil. 2020. “LAG-3 and PD-1+LAG-3 Inhibition Promote Anti-Tumor Immune Responses in Human Autologous Melanoma/T Cell Co-Cultures.”OncoImmunology 9 (1): 1736792. https://doi.org/10.1080/2162402X.2020.1736792.

Grywalska, Ewelina, Marcin Pasiarski, Stanisław Góźdź, and Jacek Roliński. 2018. “Immune-Checkpoint Inhibitors for Combating T-Cell Dysfunction in Cancer.”OncoTargets and Therapy 11: 6505–24. https://doi.org/10.2147/OTT.S150817.

Gu, Zuguang, Roland Eils, and Matthias Schlesner. 2016. “Complex Heatmaps Reveal Patterns and Correlations in Multidimensional Genomic Data.”Bioinformatics (Oxford, England) 32 (18): 2847–49. https://doi.org/10.1093/bioinformatics/btw313.

Hallek, M, K Fischer, G Fingerle-Rowson, Am Fink, R Busch, J Mayer, M Hensel, et al. 2010. “Addition of Rituximab to Fludarabine and Cyclophosphamide in Patients with Chronic Lymphocytic Leukaemia: A Randomised, Open-Label, Phase 3 Trial.”The Lancet 376 (9747): 1164–74. https://doi.org/10.1016/S0140-6736(10)61381-5.

Hamid, Omid, Henrik Schmidt, Aviram Nissan, Laura Ridolfi, Steinar Aamdal, Johan Hansson, Michele Guida, et al. 2011. “A Prospective Phase II Trial Exploring the Association between Tumor Microenvironment Biomarkers and Clinical Activity of Ipilimumab in Advanced Melanoma.”Journal of Translational Medicine 9 (November): 204. https://doi.org/10.1186/1479-5876-9-204.

Hellmann, Matthew D., Luis Paz-Ares, Reyes Bernabe Caro, Bogdan Zurawski, Sang-We Kim, Enric Carcereny Costa, Keunchil Park, et al. 2019. “Nivolumab plus Ipilimumab in Advanced Non–Small-Cell Lung Cancer.”New England Journal of Medicine 381 (21): 2020–31. https://doi.org/10.1056/NEJMoa1910231.

Hendrickx, Wouter, Ines Simeone, Samreen Anjum, Younes Mokrab, François Bertucci, Pascal Finetti, Giuseppe Curigliano, et al. 2017. “Identification of Genetic Determinants of Breast Cancer Immune Phenotypes by Integrative Genome-Scale Analysis.”OncoImmunology 0 (ja): 00–00. https://doi.org/10.1080/2162402X.2016.1253654.

Holmgaard, Rikke B., Dmitriy Zamarin, Yanyun Li, Billel Gasmi, David H. Munn, James P. Allison, Taha Merghoub, and Jedd D. Wolchok. 2015. “Tumor-Expressed IDO Recruits and Activates MDSCs in a Treg-Dependent Manner.”Cell Reports 13 (2): 412–24. https://doi.org/10.1016/j.celrep.2015.08.077.

“Https://Www.Cancer.Org/Cancer/Neuroblastoma/Detection-Diagnosis-Staging/Risk-Groups.Html.” n.d. In.

Jiang, Minlin, Keyi Jia, Lei Wang, Wei Li, Bin Chen, Yu Liu, Hao Wang, Sha Zhao, Yayi He, and Caicun Zhou. 2021. “Alterations of DNA Damage Response Pathway: Biomarker and Therapeutic Strategy for Cancer Immunotherapy.”Acta Pharmaceutica Sinica B, January, S2211383521000083. https://doi.org/10.1016/j.apsb.2021.01.003.

Kabir, Tanvir F., Aman Chauhan, Lowell Anthony, and Gerhard C. Hildebrandt. 2018. “Immune Checkpoint Inhibitors in Pediatric Solid Tumors: Status in 2018.”Ochsner Journal 18 (4): 370–76. https://doi.org/10.31486/toj.18.0055.

Keilson, Jessica M., Hannah M. Knochelmann, Chrystal M. Paulos, Ragini R. Kudchadkar, and Michael C. Lowe. 2021. “The Evolving Landscape of Immunotherapy in Solid Tumors.”Journal of Surgical Oncology 123 (3): 798–806. https://doi.org/10.1002/jso.26416.

Krijthe, Jesse H. 2015. “Rtsne: T-Distributed Stochastic Neighbor Embedding Using a Barnes-Hut Implementation.”https://github.com/jkrijthe/Rtsne.

Kudva, Anupa, and Shakeel Modak. 2019. “Immunotherapy for Neuroblastoma.” In Neuroblastoma, 147–73. Elsevier. https://doi.org/10.1016/B978-0-12-812005-7.00009-6.

Ladenstein, Ruth, Ulrike Pötschger, Dominique Valteau-Couanet, Roberto Luksch, Victoria Castel, Isaac Yaniv, Genevieve Laureys, et al. 2018. “Interleukin 2 with Anti-GD2 Antibody Ch14.18/CHO (Dinutuximab Beta) in Patients with High-Risk Neuroblastoma (HR-NBL1/SIOPEN): A Multicentre, Randomised, Phase 3 Trial.”The Lancet Oncology 19 (12): 1617–29. https://doi.org/10.1016/S1470-2045(18)30578-3.

Layer, Julian P., Marie T. Kronmüller, Thomas Quast, Debby van den Boorn-Konijnenberg, Maike Effern, Daniel Hinze, Kristina Althoff, et al. 2017. “Amplification of N-Myc Is Associated with a T-Cell-Poor Microenvironment in Metastatic Neuroblastoma Restraining Interferon Pathway Activity and Chemokine Expression.”Oncoimmunology 6 (6): e1320626. https://doi.org/10.1080/2162402X.2017.1320626.

Le, Dung T., Jennifer N. Durham, Kellie N. Smith, Hao Wang, Bjarne R. Bartlett, Laveet K. Aulakh, Steve Lu, et al. 2017. “Mismatch Repair Deficiency Predicts Response of Solid Tumors to PD-1 Blockade.”Science (New York, N.Y.) 357 (6349): 409–13. https://doi.org/10.1126/science.aan6733.

Lee, Ji Won, Meong Hi Son, Hee Won Cho, Young Eun Ma, Keon Hee Yoo, Ki Woong Sung, and Hong Hoe Koo. 2018. “Clinical Significance of *MYCN* Amplification in Patients with High-Risk Neuroblastoma.”Pediatric Blood & Cancer 65 (10): e27257. https://doi.org/10.1002/pbc.27257.

Lee, Jun Ah. 2018. “Solid Tumors in Children and Adolescents.”Journal of Korean Medical Science 33 (41): e269. https://doi.org/10.3346/jkms.2018.33.e269.

Liu, Jingkai, Qiaofei Liu, Xiang Zhang, Ming Cui, Tong Li, Yalu Zhang, and Quan Liao. 2021. “Immune Subtyping for Pancreatic Cancer with Implication in Clinical Outcomes and Improving Immunotherapy.”Cancer Cell International 21 (1): 137. https://doi.org/10.1186/s12935-021-01824-z.

Liu, Zhenqiu, Christa N. Grant, Lidan Sun, Barbara A. Miller, Vladimir S. Spiegelman, and HongGang Wang. 2020. “Expression Patterns of Immune Genes Reveal Heterogeneous Subtypes of High-Risk Neuroblastoma.”Cancers 12 (7): 1739. https://doi.org/10.3390/cancers12071739.

Lu, Rongze, Tolga Turan, Josue Samayoa, and Francesco M. Marincola. 2017. “Cancer Immune Resistance: Can Theories Converge?”Emerging Topics in Life Sciences 1 (5): 411–19. https://doi.org/10.1042/ETLS20170060.

Luke, Jason J., Riyue Bao, Randy F. Sweis, Stefani Spranger, and Thomas F. Gajewski. 2019. “WNT/β-Catenin Pathway Activation Correlates with Immune Exclusion across Human Cancers.”Clinical Cancer Research: An Official Journal of the American Association for Cancer Research, January. https://doi.org/10.1158/1078-0432.CCR-18-1942.

Moreno, Carol, Richard Greil, Fatih Demirkan, Alessandra Tedeschi, Bertrand Anz, Loree Larratt, Martin Simkovic, et al. 2019. “Ibrutinib plus Obinutuzumab versus Chlorambucil plus Obinutuzumab in First-Line Treatment of Chronic Lymphocytic Leukaemia (ILLUMINATE) : A Multicentre, Randomised, Open-Label, Phase 3 Trial.”The Lancet Oncology 20 (1): 43–56. https://doi.org/10.1016/S1470-2045(18)30788-5.

O’Neill, Rachel E., Wei Du, Hemn Mohammadpour, Emad Alqassim, Jingxin Qiu, George Chen, Philip L. McCarthy, Kelvin P. Lee, and Xuefang Cao. 2017. “T Cell-Derived CD70 Delivers an Immune Checkpoint Function in Inflammatory T Cell Responses.”Journal of Immunology (Baltimore, Md.: 1950) 199 (10): 3700–3710. https://doi.org/10.4049/jimmunol.1700380.

Perou, Charles M., Therese Sørlie, Michael B. Eisen, Matt van de Rijn, Stefanie S. Jeffrey, Christian A. Rees, Jonathan R. Pollack, et al. 2000. “Molecular Portraits of Human Breast Tumours.”Nature 406 (6797): 747–52. https://doi.org/10.1038/35021093.

Quintero Escobar, Melissa, Mariana Maschietto, Ana C. V. Krepischi, Natasa Avramovic, and Ljubica Tasic. 2019. “Insights into the Chemical Biology of Childhood Embryonal Solid Tumors by NMR-Based Metabolomics.”Biomolecules 9 (12): 843. https://doi.org/10.3390/biom9120843.

Rizvi, Naiyer A., Matthew D. Hellmann, Alexandra Snyder, Pia Kvistborg, Vladimir Makarov, Jonathan J. Havel, William Lee, et al. 2015. “Mutational Landscape Determines Sensitivity to PD-1 Blockade in Non–Small Cell Lung Cancer.”Science 348 (6230): 124–28. https://doi.org/10.1126/science.aaa1348.

Roelands, Jessica, Wouter Hendrickx, Peter J. K. Kuppen, Raghvendra Mall, Gabriele Zoppoli, Mohamad Saad, Kyle Halliwill, et al. 2019. “Genomic Landscape of Tumor-Host Interactions with Differential Prognostic and Predictive Connotations.”BioRxiv, May, 546069. https://doi.org/10.1101/546069.

Roelands, Jessica, Wouter Hendrickx, Gabriele Zoppoli, Raghvendra Mall, Mohamad Saad, Kyle Halliwill, Giuseppe Curigliano, et al. 2020. “Oncogenic States Dictate the Prognostic and Predictive Connotations of Intratumoral Immune Response.”Journal for Immunotherapy of Cancer 8 (1). https://doi.org/10.1136/jitc-2020-000617.

Rouleau, Michèle, Anand Patel, Michael J. Hendzel, Scott H. Kaufmann, and Guy G. Poirier. 2010. “PARP Inhibition: PARP1 and Beyond.”Nature Reviews Cancer 10 (4): 293–301. https://doi.org/10.1038/nrc2812.

Salerno, Elise P., Davide Bedognetti, Ileana S. Mauldin, Donna H. Deacon, Sofia M. Shea, Joel Pinczewski, Joseph M. Obeid, et al. 2016. “Human Melanomas and Ovarian Cancers Overexpressing Mechanical Barrier Molecule Genes Lack Immune Signatures and Have Increased Patient Mortality Risk.”OncoImmunology 5 (12): e1240857. https://doi.org/10.1080/2162402X.2016.1240857.

Sokol, Elizabeth, and Ami Desai. 2019. “The Evolution of Risk Classification for Neuroblastoma.”Children 6 (2): 27. https://doi.org/10.3390/children6020027.

Spranger, Stefani, and Thomas F. Gajewski. 2015. “A New Paradigm for Tumor Immune Escape: β-Catenin-Driven Immune Exclusion.”Journal for ImmunoTherapy of Cancer 3 (1): 43. https://doi.org/10.1186/s40425-015-0089-6.

Steliarova-Foucher, Eva, Murielle Colombet, Lynn A G Ries, Florencia Moreno, Anastasia Dolya, Freddie Bray, Peter Hesseling, Hee Young Shin, and Charles A Stiller. 2017. “International Incidence of Childhood Cancer, 2001–10: A Population-Based Registry Study.”The Lancet. Oncology 18 (6): 719–31. https://doi.org/10.1016/S1470-2045(17)30186-9.

Takeda, Kazuyoshi, Hideo Oshima, Yoshihiro Hayakawa, Hisaya Akiba, Machiko Atsuta, Tetsuji Kobata, Kimio Kobayashi, Mamoru Ito, Hideo Yagita, and Ko Okumura. 2000. “CD27-Mediated Activation of Murine NK Cells.”The Journal of Immunology 164 (4): 1741–45. https://doi.org/10.4049/jimmunol.164.4.1741.

Taube, J. M., A. Klein, J. R. Brahmer, H. Xu, X. Pan, J. H. Kim, L. Chen, D. M. Pardoll, S. L. Topalian, and R. A. Anders. 2014. “Association of PD-1, PD-1 Ligands, and Other Features of the Tumor Immune Microenvironment with Response to Anti-PD-1 Therapy.”Clin Cancer Res 20 (19): 5064–74. https://doi.org/10.1158/1078-0432.ccr-13-3271.

Teo, Min Yuen, Kenneth Seier, Irina Ostrovnaya, Ashley M. Regazzi, Brooke E. Kania, Meredith M. Moran, Catharine K. Cipolla, et al. 2018. “Alterations in DNA Damage Response and Repair Genes as Potential Marker of Clinical Benefit From PD-1/PD-L1 Blockade in Advanced Urothelial Cancers.”Journal of Clinical Oncology 36 (17): 1685–94. https://doi.org/10.1200/JCO.2017.75.7740.

Thorsson, Vesteinn, David L. Gibbs, Scott D. Brown, Denise Wolf, Dante S. Bortone, Tai-Hsien Ou Yang, Eduard Porta-Pardo, et al. 2018. “The Immune Landscape of Cancer.”Immunity 48 (4): 812–830.e14. https://doi.org/10.1016/j.immuni.2018.03.023.

Tumeh, Paul C., Christina L. Harview, Jennifer H. Yearley, I. Peter Shintaku, Emma J. M. Taylor, Lidia Robert, Bartosz Chmielowski, et al. 2014. “PD-1 Blockade Induces Responses by Inhibiting Adaptive Immune Resistance.”Nature 515 (7528): 568–71. https://doi.org/10.1038/nature13954.

Uyttenhove, Catherine, Luc Pilotte, Ivan Théate, Vincent Stroobant, Didier Colau, Nicolas Parmentier, Thierry Boon, and Benoît J. Van den Eynde. 2003. “Evidence for a Tumoral Immune Resistance Mechanism Based on Tryptophan Degradation by Indoleamine 2,3-Dioxygenase.”Nature Medicine 9 (10): 1269–74. https://doi.org/10.1038/nm934.

Valentijn, L. J., J. Koster, F. Haneveld, R. A. Aissa, P. van Sluis, M. E. C. Broekmans, J. J. Molenaar, J. van Nes, and R. Versteeg. 2012. “Functional MYCN Signature Predicts Outcome of Neuroblastoma Irrespective of MYCN Amplification.”Proceedings of the National Academy of Sciences 109 (47): 19190–95. https://doi.org/10.1073/pnas.1208215109.

Wei, Jun S., Igor B. Kuznetsov, Shile Zhang, Young K. Song, Shahab Asgharzadeh, Sivasish Sindiri, Xinyu Wen, et al. 2018. “Clinically Relevant Cytotoxic Immune Cell Signatures and Clonal Expansion of T Cell Receptors in High-Risk MYCN-Not-Amplified Human Neuroblastoma.”Clinical Cancer Research : An Official Journal of the American Association for Cancer Research 24 (22): 5673–84. https://doi.org/10.1158/1078-0432.CCR-18-0599.

Wilkerson, Matthew D., and D. Neil Hayes. 2010. “ConsensusClusterPlus: A Class Discovery Tool with Confidence Assessments and Item Tracking.”Bioinformatics 26 (12): 1572–73. https://doi.org/10.1093/bioinformatics/btq170.

Wolf, A. 2004. “Overexpression of Indoleamine 2,3-Dioxygenase in Human Inflammatory Bowel Disease.”Clinical Immunology 113 (1): 47–55. https://doi.org/10.1016/j.clim.2004.05.004.

Xie, Xiaoliang, Yumei Li, Haixia Zhu, Zhixing Kuang, Deta Chen, and Tianyou Fan. 2020. “Prognostic Significance of β-Catenin Expression in Osteosarcoma: A Meta-Analysis.”Frontiers in Oncology 10: 402. https://doi.org/10.3389/fonc.2020.00402.

Yu, Alice L., Andrew L. Gilman, M. Fevzi Ozkaynak, Wendy B. London, Susan G. Kreissman, Helen X. Chen, Malcolm Smith, et al. 2010. “Anti-GD2 Antibody with GM-CSF, Interleukin-2, and Isotretinoin for Neuroblastoma.”The New England Journal of Medicine 363 (14): 1324–34. https://doi.org/10.1056/NEJMoa0911123.

Yust-Katz, Shlomit, Simon Khagi, and Mark R. Gilbert. 2020. “Neurologic Complications.” In Abeloff’s Clinical Oncology, 688–706.e7. Elsevier. https://doi.org/10.1016/B978-0-323-47674-4.00045-1.

Zhang, Chi, Jing-Hui Zheng, Zong-Han Lin, Hao-Yuan Lv, Zhuo-Miao Ye, Yue-Ping Chen, and Xiao-Yun Zhang. 2020. “Profiles of Immune Cell Infiltration and Immune-Related Genes in the Tumor Microenvironment of Osteosarcoma.”Aging 12 (4): 3486–3501. https://doi.org/10.18632/aging.102824.

Zhang, Jing, David J. H. Shih, and Shiaw-Yih Lin. 2020. “Role of DNA Repair Defects in Predicting Immunotherapy Response.”Biomarker Research 8 (1): 23. https://doi.org/10.1186/s40364-020-00202-7.

Zhang, Peng, Xiaofang Wu, Moushumi Basu, Chen Dong, Pan Zheng, Yang Liu, and Anthony David Sandler. 2017. “MYCN Amplification Is Associated with Repressed Cellular Immunity in Neuroblastoma: An In Silico Immunological Analysis of TARGET Database.”Frontiers in Immunology 8 (November): 1473. https://doi.org/10.3389/fimmu.2017.01473.

Zhang, Ya, and Xin Wang. 2020. “Targeting the Wnt/β-Catenin Signaling Pathway in Cancer.”Journal of Hematology & Oncology 13 (1): 165. https://doi.org/10.1186/s13045-020-00990-3.

