## Supplementary materials for "The Immune landscape of pediatric solid tumors"

Supplementary Figure1 . Pearson correlation of ICR genes in different pediatric tumors

**A.**

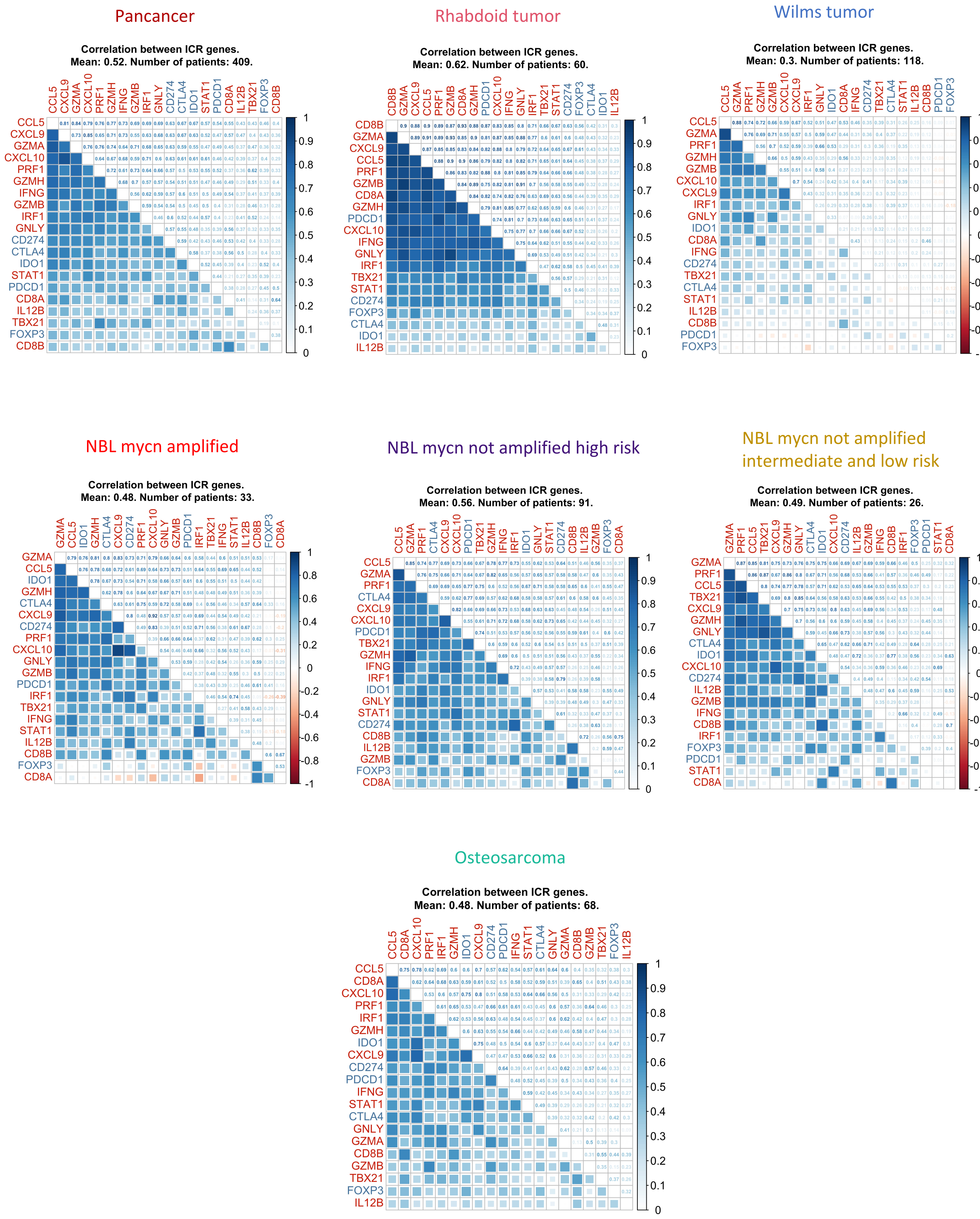

Supplementary Figure 2. The Immunologic constant of rejection

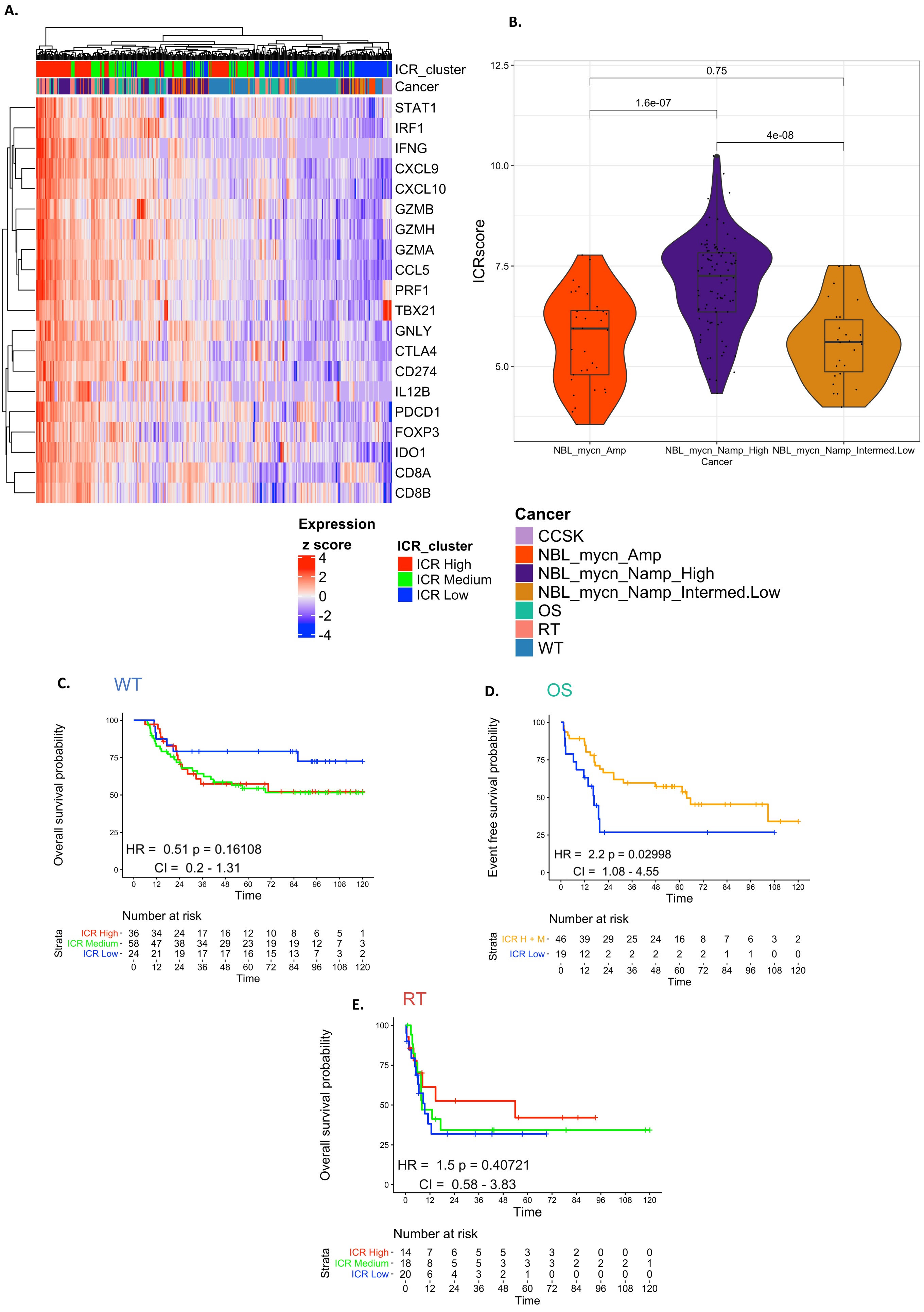

Supplementary Figure 3. Intrinsic oncogenic pathways across pediatric tumors

A.

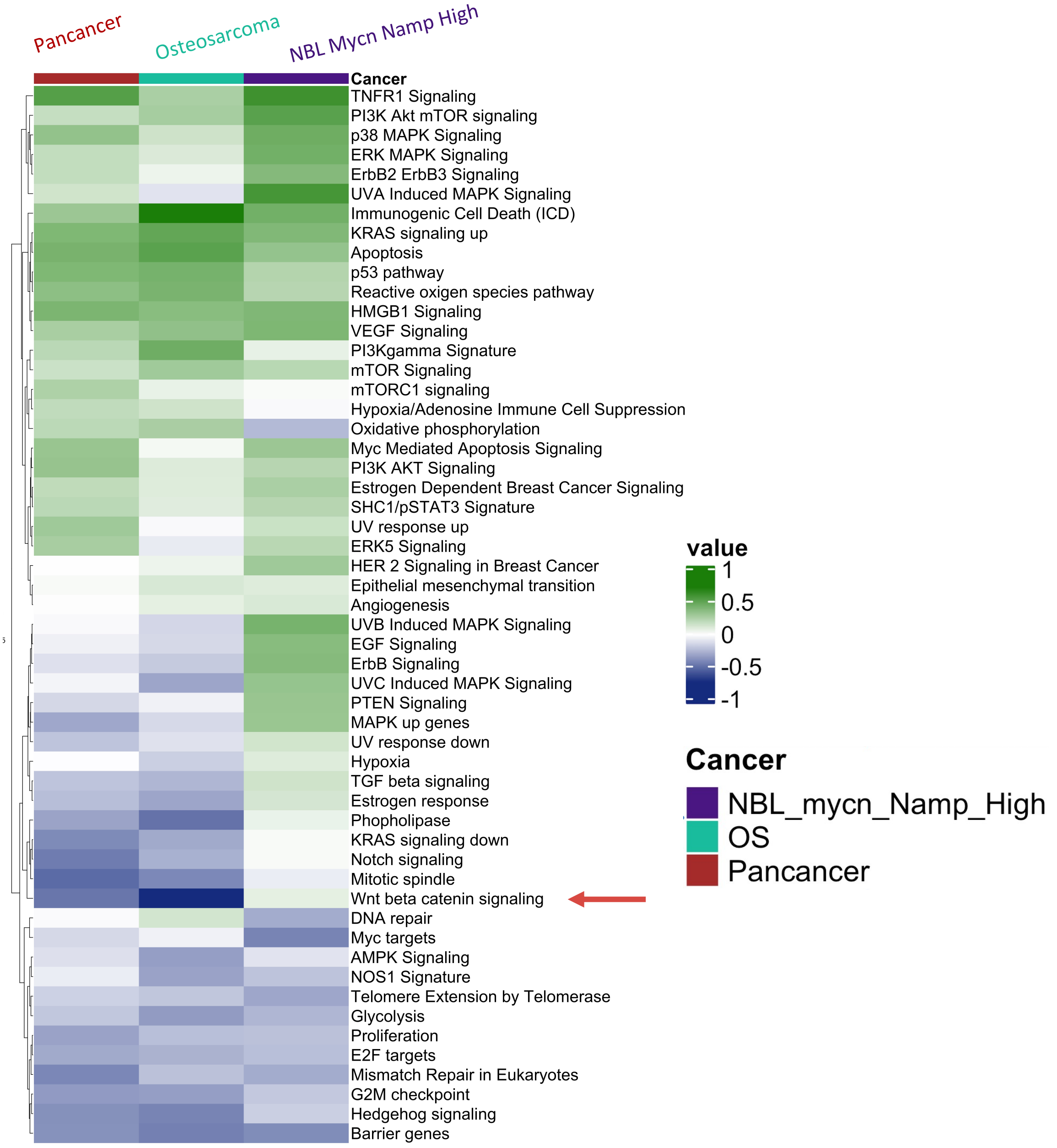


B.

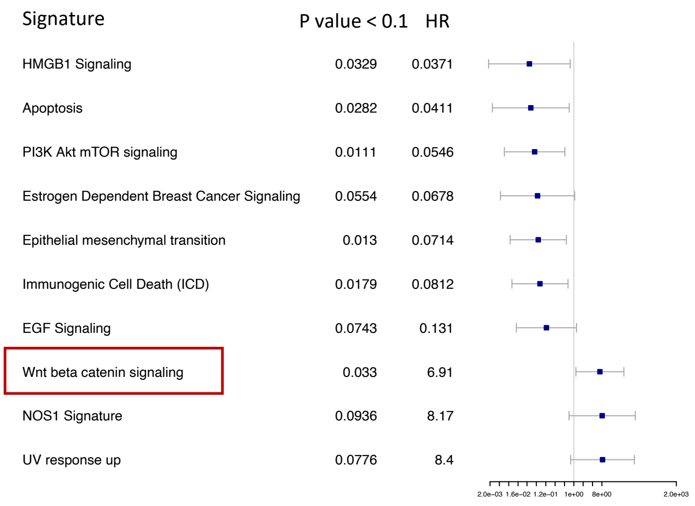


Osteosarcoma

C.

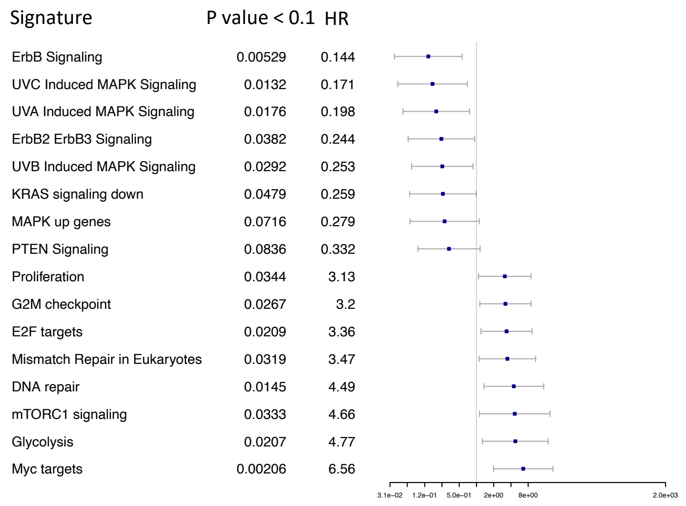


High risk MYCN Not amplified NBL

Supplementary Figure 4. Overall survival across Immune subtypes within cancer types

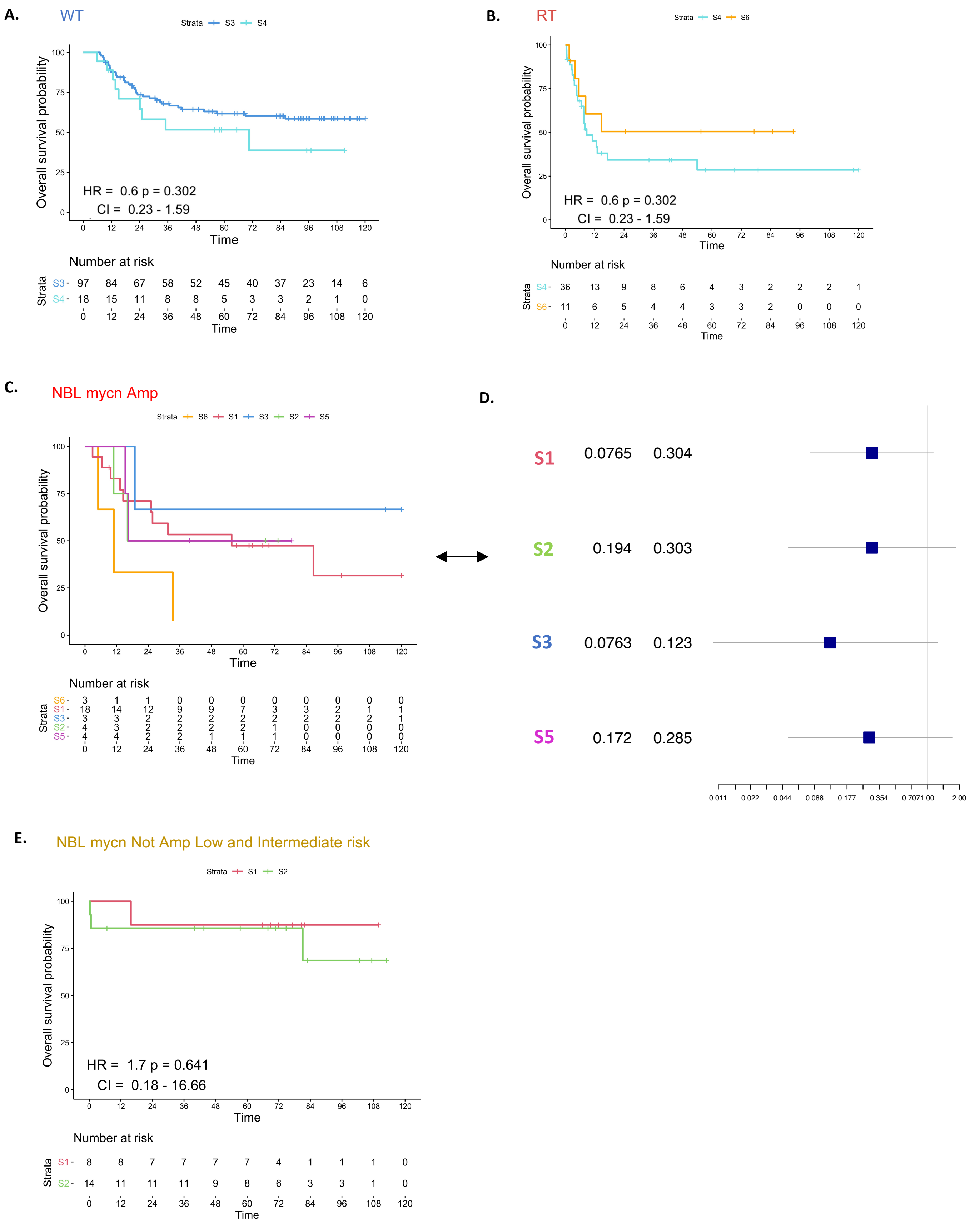

Supplementary Figure 5. CIBERSORTx immune cells proportions across Immune subtypes

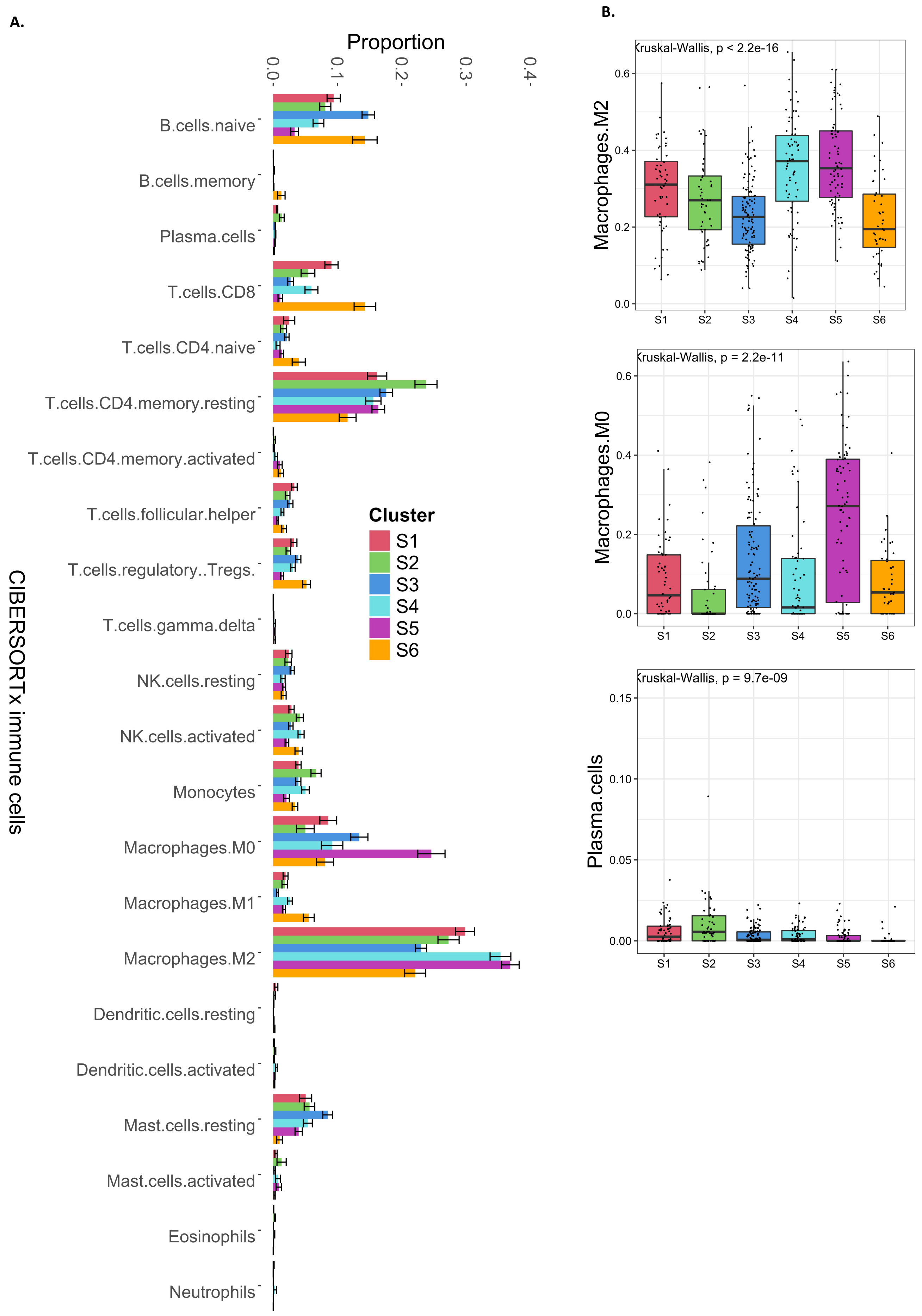

Supplementary Figure 6. CIBERSORTx immune cells proportions across Immune subtypes (Aggregate)

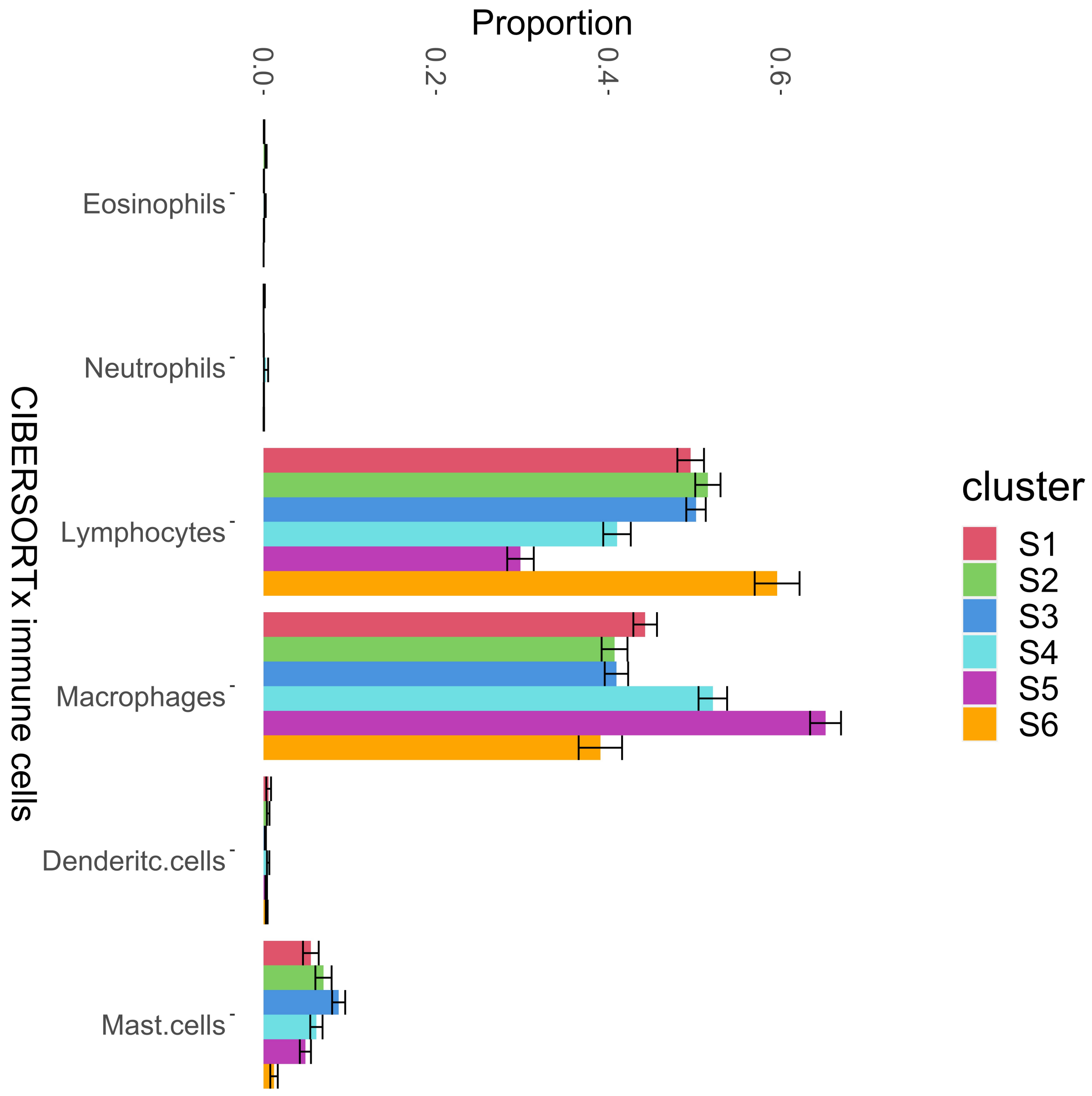
